# Mono-Ubiquitylation-Dependent Rap2 Activation Regulates Lamellipodia Dynamics During Cell Migration

**DOI:** 10.1101/2025.04.25.650668

**Authors:** Andrew Neumann, Revathi Sampath, Emily Mayerhofer, Valeryia Mikalayeva, Vytenis Arvydas Skeberdis, Ieva Sarapinienė, Rytis Prekeris

**Affiliations:** Department of Cell and Developmental Biology, School of Medicine, University of Colorado Anschutz Medical Campus, Aurora, CO 80045, USA; Lithuanian University of Health Sciences, Kaunas, Lithuania

## Abstract

Cell migration is a complex process hallmarked by front-to-back cell polarity that is established by the highly dynamic actin cytoskeleton. Branched actin polymerization creates a lamellipodia at the leading edge of the cell, while the contractile acto-myosin cytoskeleton is present at the lagging edge. Rap2, a Ras GTPase family member, has previously been reported to localize to the lamellipodia as a result of Rab40/CRL5 E3 ubiquitin ligase induced tri-mono-ubiquitylation. However, how Rap2 functions and how mono-ubiquitylation targets Rap2 to the lamellipodia remained unclear. Here, we demonstrate that Rap2 is recruited to retracting lamellipodia ruffles where it inhibits RhoA and regulates lamellipodia dynamics and facilitates cell migration. Furthermore, using a variety of genetic and pharmacological techniques, we show that tri-mono-ubiquitylation is required for GEF-dependent Rap2 activation, a necessary step for Rap2 targeting to lamellipodia membrane. As such, we demonstrate how this unique mono-ubiquitylation of Rap2 regulates lamellipodia actin dynamics during cell migration.

**SUMMARY:** Mono-ubiquitylation is necessary for activity and localization of the Rap2 GTPase. Here, we report that Rap2 lamellipodia and membrane dwell time is facilitated by mono-ubiquitylation dependent GEF activation. Furthermore, we demonstrate that mono-ubiquitylated Rap2 regulates lamellipodia dynamics through inhibiting RhoA activity.

## INTRODUCTION

Cell migration is a complex process essential for multiple functions including organism development and immune responses (Bravo-Cordero et al., 2012; Franz et al., 2002; Trepat et al., 2012). A plethora of factors must be tightly coordinated for proper cell migration. One of which, the actin cytoskeleton, is essential for providing the dynamic structures necessary for cellular locomotion (Neumann and Prekeris, 2023; Schaks et al., 2019). The actin cytoskeleton polymerizes to form multiple uniquely constructed structures which, in a migrating cell, are spatially segregated to create front-to-back polarity. Simply put, the front, or leading edge, of the cell is characterized by Arp2/3-dependent branched actin filaments which push the front of the cell forward, creating a ruffling structure known as the lamellipodia. The rear, or lagging edge, is canonically defined by focal adhesion attached linear actin filaments connected by non-muscle myosin II motors (NMII) to form acto-myosin stress fibers. When activated, NMII facilitates acto-myosin stress fiber contraction which pulls the lagging edge forward (Bisi et al., 2013; Fregoso et al., 2022; Hotulainen and Lappalainen, 2006; Kolega, 2006; Mullins et al., 1998; Naumanen et al., 2008; Suraneni et al., 2012). These distinct actin structures must be tightly spatiotemporally regulated to maintain cell movement in the appropriate direction.

Regulation of the actin cytoskeleton is largely facilitated by small monomeric GTPases. GTPases act as molecular switches that cycle between an active, GTP-bound, and an inactive, GDP-bound state. Activation of GTPases is facilitated by Guanine Nucleotide Exchange Factors (GEFs) and inactivation by GTPase Activating Proteins (GAPs) (Bos et al., 2007). The Rho family of GTPases is largely accepted as the master regulators of actin during cell migration.

Among several Rho GTPases, Rac1 is generally considered to be the major regulator of leading edge actin dynamics. Rac1 activates the Arp2/3 complex, resulting in branched actin polymerization and lamellipodia formation (Gorelik and Gautreau, 2015; Kraynov et al., 2000; Kurokawa et al., 2003; Molinie and Gautreau, 2017; Simanov et al., 2021; Suraneni et al., 2012). Oppositely, RhoA is accepted as the main regulator of lagging edge actin structure by activating its effectors which stimulate NMII activity, resulting in acto-myosin stress fiber contraction (Hotulainen and Lappalainen, 2006; Julian and Olson, 2014; Kolega, 2006; Kurokawa and Matsuda, 2005; Naumanen et al., 2008; Watanabe et al., 1999; Wong et al., 2023). Precise coordination of these major events of leading edge extension and lagging edge retraction provide the basic mechanics for cell locomotion. Thus, regulation of RhoA and Rac1 localization and activation through GEFs and GAPs has become a major focus of study in the field of cell migration.

Despite the importance of Rho GTPases, other GTPases have also been shown to regulate actin dynamics associated with cell migration. Specifically, the Ras family of GTPases is known for regulating actin cytoskeleton dynamics and cell migration, largely through the Ras subfamily of GTPases (HRAS, KRAS, NRAS) which have been found to be mutated in patients and drive cancer metastasis (Bos et al., 2007; Collins et al., 2023; Fuentes-Calvo et al., 2013; Prior et al., 2012). Less commonly studied among the Ras GTPases is the Rap GTPase subfamily (Rap1a, Rap1b, Rap2a, Rap2b, Rap2c). Rap1 and Rap2 have been shown to regulate endothelial barriers through indirectly regulating RhoA. However, the function of these proteins remains less understood in other cellular contexts such as cell migration (Pannekoek et al., 2013; Post et al., 2015). Rap1 has been previously implicated in regulating cell migration by mediating focal adhesion formation through integrin trafficking (Reedquist et al., 2000; Rothenberg et al., 2023). Nevertheless, while we have previously shown the importance of Rap2 in cell motility (Duncan et al., 2022), its function and regulation during cell migration remains largely unclear.

Our previous work has shown that Rap2 is a substrate of a Rab40/Cullin5 E3 ubiquitin ligase complex (Rab40/CRL5), and that Rap2 is - mono-ubiquitinated at K117, K148, and K150. These modifications are necessary for Rap2 to be targeted to the lamellipodia plasma membrane. Furthermore, mono-ubiquitylation facilitates Rap2 to be in its active, GTP-bound state. Inhibition of Rab40/CRL5-dependent mono-ubiquitylation of Rap2 results in inactivation of Rap2 molecules that accumulate in and are degraded by lysosomes (Duncan et al., 2022).

Results from this study have shown that Rap2 is dynamically regulated in a unique way which ascribed the need for further study into Rap2 regulation and function. Specifically, we sought to understand the question of how Rap2 mono-ubiquitylation facilitates its conversion to a GTP-bound (active) state, how mono-ubiquitylation leads to Rap2 targeting to the lamellipodia plasma membrane, and how ubiquitylated Rap2 affects cell migration.

Here we demonstrate that Rap2 facilitates cell migration by inhibiting RhoA at the lamellipodia, thus contributing to the establishment of front-to-back polarity in migrating cells. Importantly, we demonstrate that Rap2 is recruited to the lamellipodia plasma membrane during lamellipodia ruffle retraction where it appears to inhibit RhoA. Thus, Rap2 recruitment to the retracting lamellipodia limits the extent of ruffle retraction and regulates lamellipodia dynamics and cell migration. We also show that mono-ubiquitylation of Rap2 is essential for its function in regulating RhoA activity and its downstream functions during migration. Furthermore, using a combination of techniques to inhibit membrane trafficking and alter Rap2 activity, we demonstrate that mono-ubiquitylation of Rap2 at K117, K148, and K150 is necessary for Rap2 activation by its GEF, which increases Rap2 activation and dwell-time at the lamellipodia plasma membrane. Accordingly, we propose a model in which the Rap2 GTPases are regulated in their migratory functions through mono-ubiquitylation by the Rab40/CRL5 complex. Ubiquitylated Rap2 is activated, facilitating its retention at the lamellipodia plasma membrane where it regulates leading edge dynamics by inhibiting RhoA-driven retraction of actin ruffles.

## RESULTS

### Rap2 is required for cell migration and lamellipodia dynamics

While Rap2 has been previously shown to be essential for MDA-MB-231 cell migration and invasion (Duncan et al., 2022), how and when Rap2 regulates membrane and actin dynamics at the lamellipodia in migrating cells remains to be fully understood. To determine the molecular machinery governing the role of Rap2 during migration we created a Rap2a, Rap2b, and Rap2c co-knock-out MDA-MB-231 (Rap2-KO) line using CRISPR/Cas9 (Duncan et al., 2022). In order to better understand what aspect of cell migration is mediated by Rap2, we performed individual cell random migration assays using time-lapse imaging. Consistent with the involvement of Rap2 in regulating cell migration, Rap2-KO cells moved slower, and a shorter distance as compared to control cells (Fig.1A-C, Video 1 and Video 2). Because Rap2 has been shown to be enriched at the lamellipodia, we next assessed whether changes in lamellipodia dynamics contributed to the observed migratory defects.

**Figure 1:**
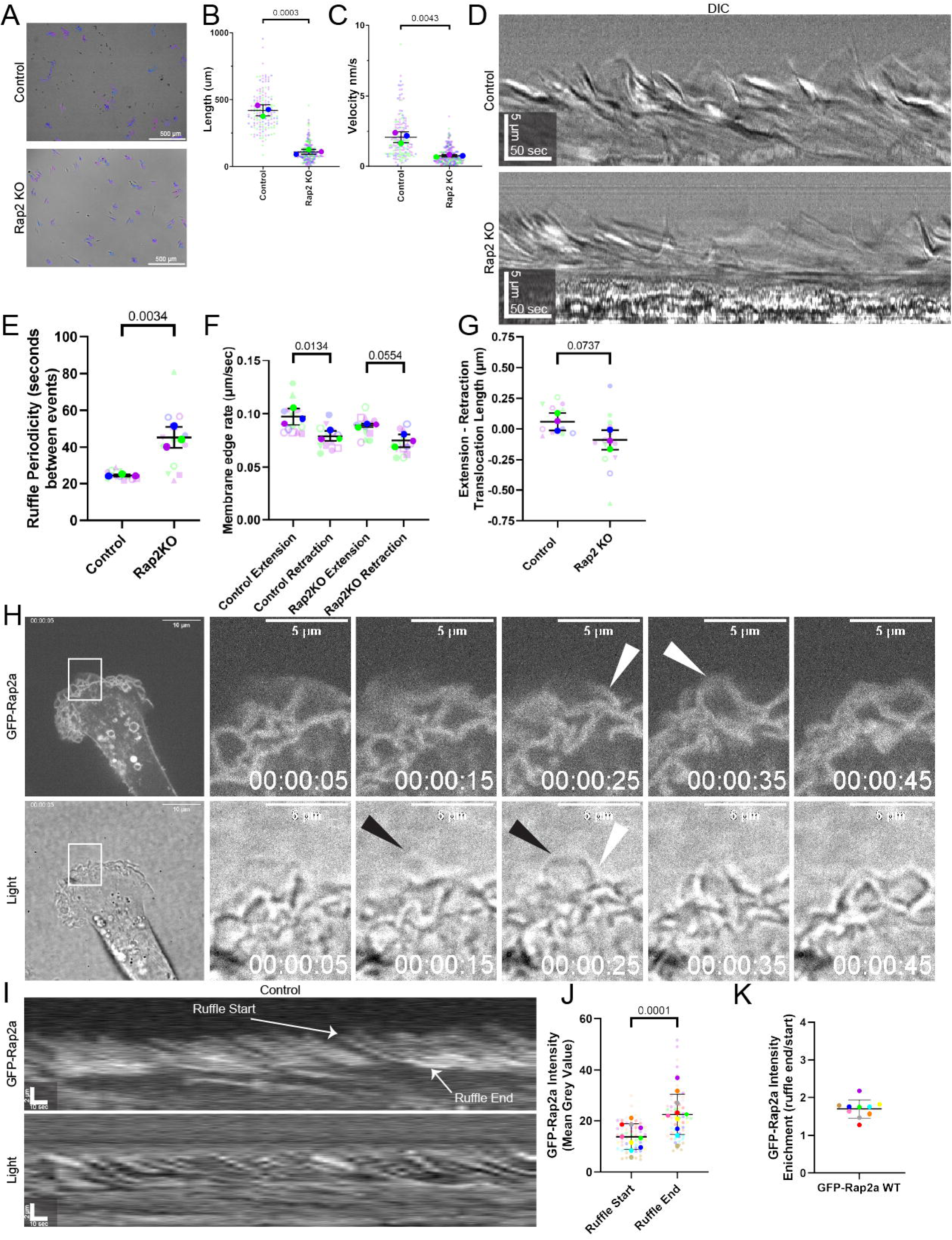
Rap2 is required for cell migration and lamellipodia dynamics. **(A-C)** Tracking of control and Rap2KO cells from time-lapse imaging of individual cell random migration assays (Videos 1 and 2). Quantifications show the total length of the track the cells migrated **(B)** and velocity at which the cells migrated **(C)**. **(D-G)** Kymographs taken from SACED time-lapse imaging of control and Rap2KO cells (Videos 3 and 4). Actin ruffle periodicity **(E)** and membrane extension and retraction length **(F-G)** were measured from SACED kymographs. **(H)** GFP-Rap2 expressing MDA-MB-231 cells were analyzed by time-lapse microscopy (see Video 5). Shown are selected GFP-Rap2a and transmitted light images taken from time-lapse series. Black arrows point to an extending ruffle that lacks GFP-Rap2a. White arrow points to retracting ruffle with accumulating GFP-Rap2a. **(I-K)** Kymographs generated from GFP-Rap2a time-lapse series (Video 5). Ruffle start and ruffle end are labeled as examples of where mean grey value was measured. **(J, K)** Quantification of mean grey value from the start and end of ruffles from kymographs of GFP-Rap2a dynamics showing that GFP-Rap2a accumulates during ruffle retraction.

Stroboscopic Analysis of Cell Dynamics (SACED) was performed using previously established calculations from kymographs generated from time-lapse microscopy of migrating MDA-MB-231 cells, allowing for the analysis of lamellipodia ruffling and extension/retraction dynamics ((Hinz et al., 1999), Fig.1D, Fig. S1A, Video 3 and Video 4). Noticeably, while Rap2 KO had little effect on ruffle extension and retraction rates (Fig. S1B-D) it significantly decreased ruffle periodicity as compared to control cells (Fig. 1E). Rap2-KO cells also exhibited increased membrane retraction distance (Fig. 1F). This increased retraction distance of lamellipodia ruffles in Rap2-KO cells resulted in lamellipodia that retract more than they extend (Fig. 1G), likely contributing to the inability for Rap2-KO cells to move. In accordance with increased retraction, in all our time-lapse videos Rap2-KO cells were consistently seen to have large retraction events occur at the lamellipodia during ruffling (Video 4). In contrast, control cells exhibited constantly ruffling lamellipodia with membranes that extend a longer distance than they retract, thus contributing to forward cell movement (Fig. 1G).

To understand how Rap2 regulates lamellipodia extension/retraction dynamics we generated an MDA-MB-231 cell line stably expressing GFP-Rap2a-WT (wild type) and analyzed the dynamics of GFP-Rap2-WT recruitment to the lamellipodia using time-lapse microscopy (Video 5). As shown in the figure 1H, GFP-Rap2a-WT is largely absent at the lamellipodia as the membrane extends forward (black arrowheads) and accumulates on the ruffle as it retracts (white arrowheads). Interestingly, the fluorescent intensity of GFP-Rap2a-WT increases as the ruffle retracts, indicating an increasing accumulation of Rap2a throughout ruffle retraction (Fig1. I-K). This increase in Rap2 localization in the retracting lamellipodia suggests that Rap2 may function to terminate lamellipodia ruffle retraction. This hypothesis is consistent with the data that Rap2 KO leads to an increase in lamellipodia retraction distance and a decrease in lamellipodia ruffling periodicity (Fig. 1E-G).

Previous data suggest that in endothelial cells, Rap GTPases can indirectly modulate RhoA activity. Thus, we next asked if RhoA is also recruited to retracting ruffles and whether Rap2 may regulate lamellipodia ruffling through modulation of RhoA activity. Accordingly, we performed RhoA activity pulldowns using GST-RBD (Rhotekin binding domain, a known RhoA effector that binds to active RhoA) beads. More RhoA was found to pulldown in lysates from Rap2-KO cells as compared to control cells (Fig. S1E-F), suggesting that Rap2 negatively regulates RhoA. To test whether Rap2 may inhibit RhoA at the retracting ruffles, we co-transfected MDA-MB-231 cells with GFP-Rap2a and activated RhoA Biosensor (dTom-2xrGBD, (Mahlandt et al., 2021)). Consistent with its role in driving ruffle retraction, activated RhoA can be observed to quickly accumulate at the initiation of lamellipodia ruffle retraction where it colocalizes with GFP-Rap2a (Fig. 2A-B). Furthermore, as GFP-Rap2 levels increase towards the termination of ruffle retraction, the levels of activated RhoA begin to plateau then decrease (Fig. 2C). Taken together, these data indicate that Rap2 plays a role in regulating lamellipodia dynamics and cell migration by regulating the extent of the lamellipodia retraction through inhibition RhoA.

**Figure 2:**
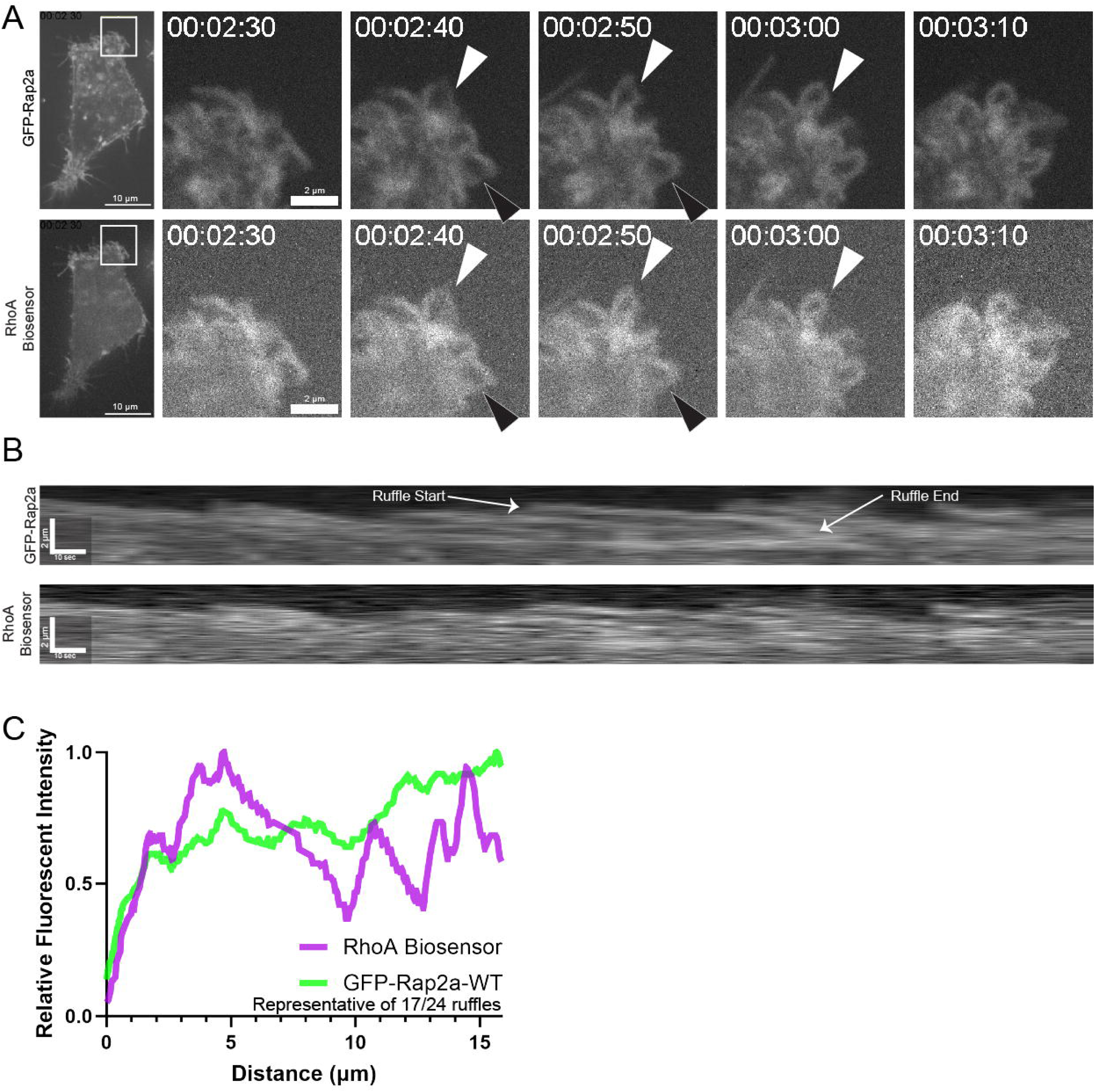
Rap2 and active RhoA colocalize in retracting lamellipodia ruffles. **(A)** Time-lapse analysis of MDA-MB-231 cells co-expressing GFP-Rap2a and dTom-2xrGBD (RhoA biosensor) (also see Video 6). Shown are selected GFP-Rap2a and RhoA biosensor images taken from the time-lapse series of a ruffling lamellipodia. White arrows show a ruffle where GFP-Rap2a is recruited with the RhoA biosensor. Black arrows show ruffles where GFP-Rap2a recruitment decreases fluorescent intensity of the RhoA biosensor. **(B-C)** Kymographs generated from GFP-Rap2a and dTom-2xrGBD time-lapse series (Video 6). White ruffle outline is a representation of where ruffle fluorescent intensity was measured. **(C)** Line scan analysis of relative GFP-Rap2a and dTom-2xrGBD fluorescent intensities as measured from the ruffle outlined in **(B).** Of the 24 ruffles analyzed (3 ruffles/cell from 8 different cells), 17 displayed a pattern as in **(C)** where dTom-2xrGBD signal either decreased or plateaued as GFP-Rap2a intensity increased.

### Rap2 is required for establishing front-to-back polarity in migrating cells

The Rap2 KO induced defects in lamellipodia extension/retraction suggest that Rap2 regulates leading edge dynamics which are necessary for the establishment of front-to-back polarity in migrating cells. To test whether Rap2 is also required for the establishment of front-to-back polarity, we used immunofluorescence microscopy (IF) to visualize the actin cytoskeleton in Rap2-KO cells. As previously shown, MDA-MB-231 cells exhibit a front-to-back polarization hallmarked by a wide leading lamellipodia with clearly identifiable actin ruffles and a discrete and narrow lagging edge (Fig. 3A). In contrast, Rap2-KO cells did not have clear leading or lagging edges and exhibited a visible increase in stress fibers and surface area (Fig. 3A-B). Cell aspect ratio can be used as a measurement to analyze front-to-back polarization (Wong et al., 2023). Consistent with the role of Rap2 in the establishment of front-to-back polarity, Rap2 KO led to a decrease in aspect ratio (Fig. 3C).

**Figure 3:**
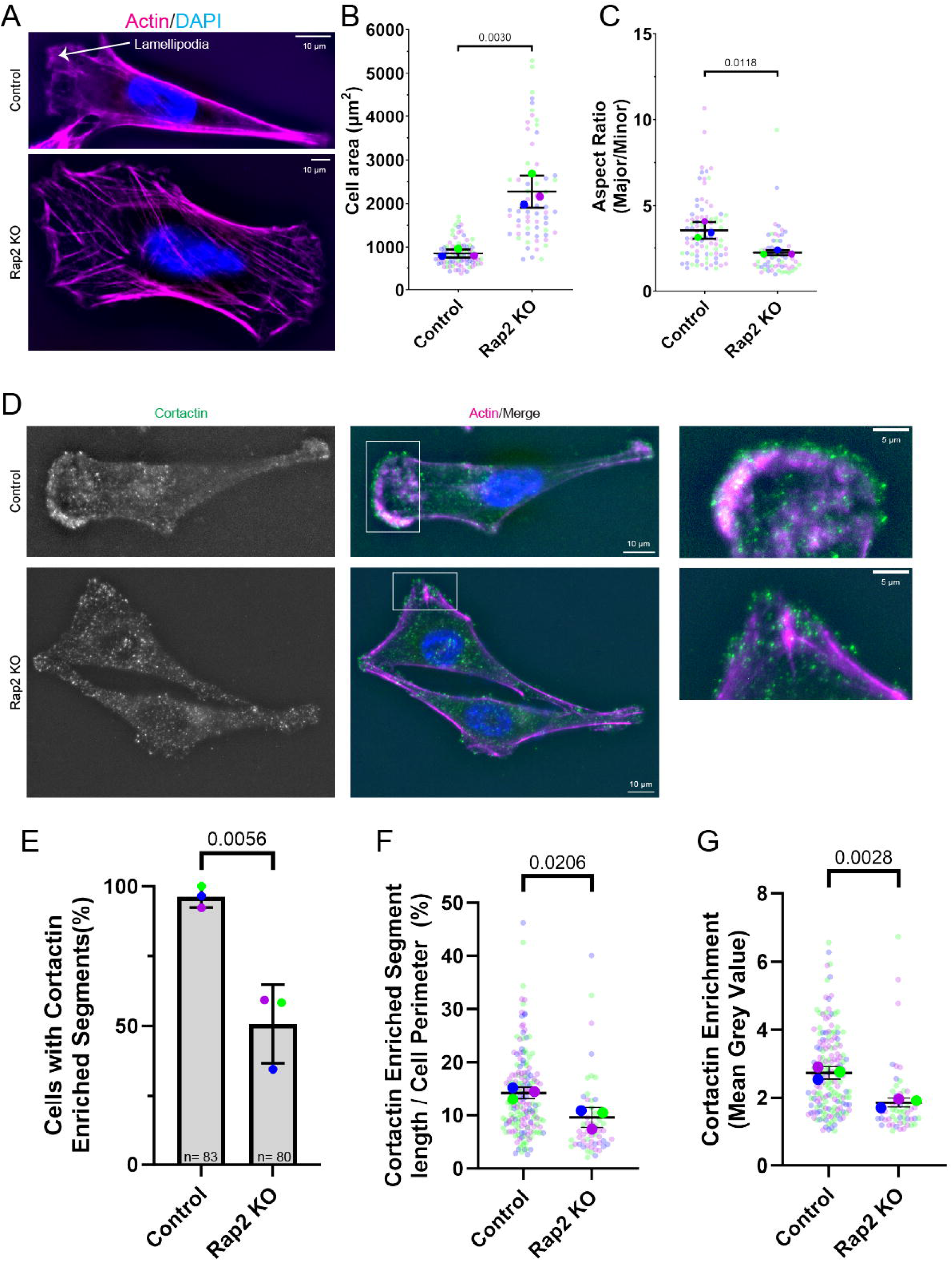
Rap2 regulates the formation of front-to-back polarity in migrating cells. **(A-C)** Phalloidin-Alexa596 staining of Rap2-KO and control MDA-MB-231 cells to visualize the actin cytoskeleton. **(B-C)** Quantification of cell size **(B)** and aspect ratio **(C)**. **(D)** Control and Rap2-KO MDA-MB-231 cells were fixed and stained with phalloidin-Alexa596 and an anti-cortactin (marker for dynamic lamellipodia) antibody. **(E-G)** Quantification of anti-cortactin antibody signal in MDA-MB-231 cells. Panel **(E)** shows the percentage of cells with cortactin enriched segments. Panel **(F)** shows the percentage of the individual cell perimeter made up of a cortactin enriched segments. Panel **(G)** shows the enrichment of cortactin in the lamellipodia as compared to the actin cortex.

Loss of front-to-back polarity and perturbed lamellipodia dynamics indicate that Rap2 depletion may also affect the formation of the lamellipodia. To assess the potential of Rap2-KO cell to make a dynamic lamellipodia, we used cortactin, an actin-binding protein enriched at areas of Arp2/3-dependent branched actin polymerization, as a molecular marker of dynamic lamellipodia (Kaksonen et al., 2000; Weed et al., 2000). As shown in figure 3D-E, only around 50% of Rap2-KO cells had lamellipodia-like structures with visible cortactin enrichment, indicating that most Rap2-KO cells are incapable of creating lamellipodia. Of the population of Rap2-KO cells with visible cortactin-enriched domains, the lamellipodia-like structures were significantly smaller and displayed less cortactin enrichment as compared to the controls (Fig. 3F-G). Thus, while some Rap2-KO cells can form lamellipodia-like structures, these lamellipodia are smaller and likely contain less dynamic branched actin, further explaining defects in lamellipodia dynamics (Fig. 1D-G). Accordingly, loss of Rap2 impacts the formation of critical migratory structures such as lamellipodia and ablates front-to-back polarity in migrating cells.

### Loss of Rap2 leads to increased formation of FAs and acto-myosin stress fibers

Our data demonstrate that Rap2 is involved in regulating cell polarity and migration in part through regulation of RhoA activity during lamellipodia extension/retraction. However, upon Rap2 KO, we noticed morphological defects throughout the whole cell which may originate as a result of an improper increase and propagation of RhoA activity. One of the well-established functions of RhoA is to regulate stress fiber tension and Focal Adhesion (FA) formation (Julian and Olson, 2014; Mishra and Manavathi, 2021; Wolfenson et al., 2011). Thus, we next investigated whether Rap2 regulates FA formation and dynamics throughout the entire cell. To that end, we transfected control and Rap2-KO cells with GFP-Paxillin and analyzed FA dynamics using time-lapse microscopy. As expected, migrating control MDA-MB-231 cells exhibit dynamic nascent FAs, characterized by rapid turnover at the lamellipodia, and cytoplasmic FAs with relatively short lifespans and rapid movement across the dorsal cell surface during cell migration (Fig. 4A-D, Video 6). In contrast, FAs in Rap2-KO cells did not turn over at lamellipodia-like structures, lasted throughout our whole time-lapse series, and moved slowly across the dorsal cell surface (Fig. 4A-D, Video 7). These results are consistent with FAs in Rap2-KO cells having increased stability and having disassembly defects.

**Figure 4:**
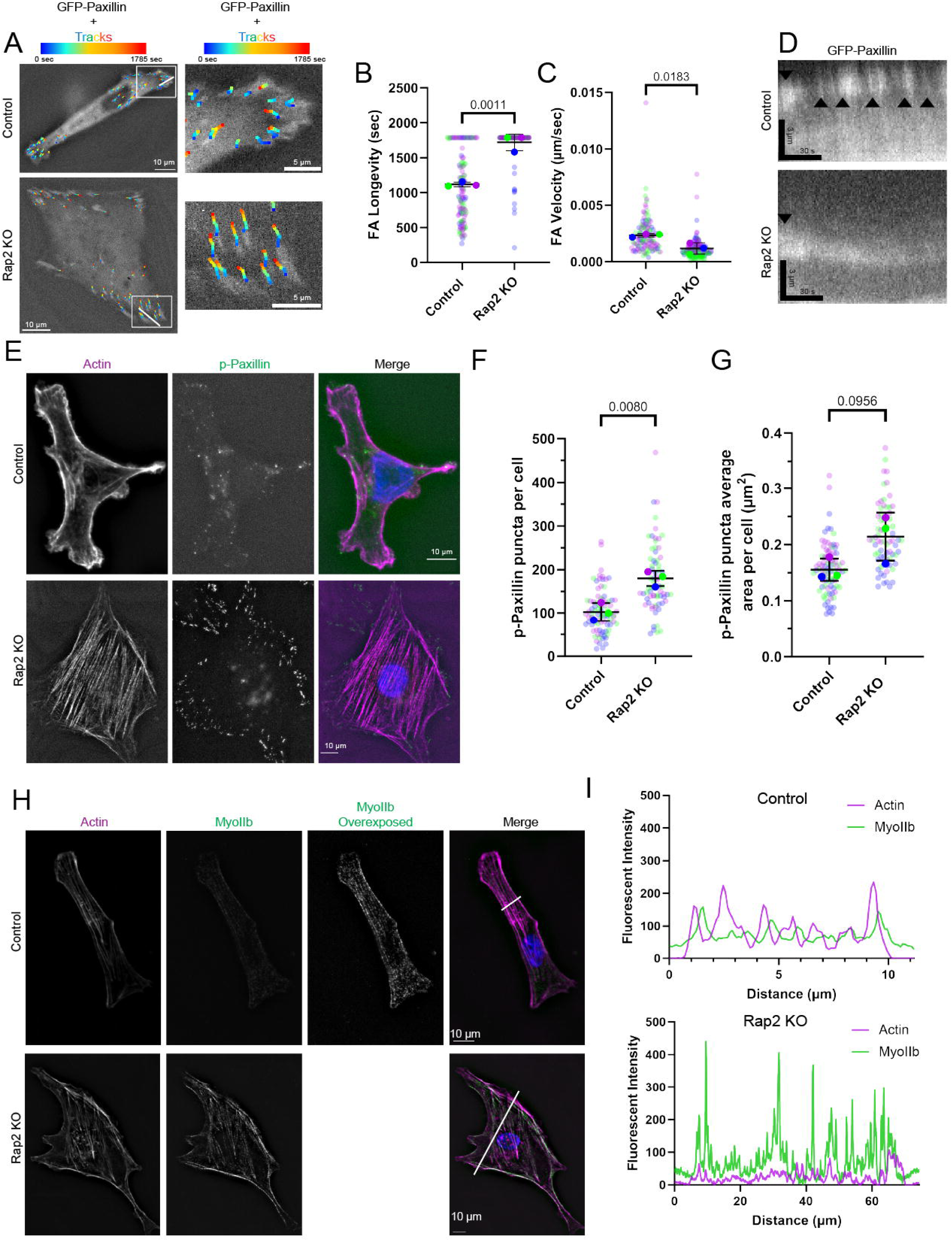
Rap2 regulates the formation and dynamics of Focal Adhesions. **(A-D)** Time-lapse analysis of MDA-MB-231 cells expressing GFP-Paxillin. Tracks of paxillin puncta movement over time are overlayed on top of the first image taken. **(B-C)** Quantification of paxillin puncta longevity **(B)** and velocity **(C)**. Panel (**D)** shows kymographs generated from GFP-Paxillin time-lapse series (Videos 7-8). Kymographs were generated from the line drawn in panel **(A)**. To visualize the spatiotemporal dynamics of individual focal adhesions, line was drawn through a focal adhesion shown in panel **(A)**. Black arrows mark the appearance of new focal adhesion as marked by GFP-Paxillin signal. **(E-G)** Control and Rap2-KO MDA-MB-231 cells were stained with phalloidin-Alexa596 (Actin) and an anti-phosphorylated(p)-Paxillin antibody. **(F-G)** Quantification of focal adhesions from cells shown in panel **(E)**. Panel **(F)** shows the average number of p-Paxillin puncta per cell and panel **(G)** shows the average size of p-Paxillin puncta per cell. **(H-I)** Control and Rap2-KO MDA-MB-231 cells were stained with phalloidin-Alexa596 (Actin) and an anti-MyoIIb antibody. Panel **(I)** shows line scans taken from lines drawn across images shown in panel **(H)**.

To further analyze the FA phenotypes observed in our time-lapse imaging, we performed IF to visualize endogenous FAs using an anti-phospho-paxillin antibody (Bachmann et al., 2022). In control cells phospho-paxillin labeled FAs, localized predominantly in the cell periphery, with FA clusters noted at areas of actin ruffling (Fig. 4E). However, in Rap2-KO cells, phospho-paxillin labelled FAs were not only in the cell periphery but could also be observed towards the cell center. Furthermore, in Rap2-KO cells, there were more phospho-paxillin labelled FAs and FAs were bigger than in control cells, further suggesting an increase in FA stability (Fig. 4E-G).

FAs are involved in mediating cytoskeletal tension and allow for the production of contractile forces generated by NMII motors that assemble on FA-anchored stress fibers (Kolega, 2006; Richter et al., 2021). Accordingly, we predicted that the increase in FAs number and size in Rap2-KO cells correlated with an increase in contractility of acto-myosin stress fibers. To test this hypothesis, we performed IF microscopy using an anti-NMIIb antibody and analyzed NMIIb distribution along actin stress fibers. As shown in the figure 4H-I, Rap2-KO cells had increased NMIIb fluorescent intensity that colocalized with stress fibers. Thus, our observations regarding FAs in Rap2-KO cells are linked with changes in actin-myosin stress fiber contractility. As downstream effectors of RhoA, increased FA stability and acto-myosin contractility concur with our data suggesting Rap2 negatively regulates RhoA (Fig. 2, Fig. S1E-F).

### Rap2 mono-ubiquitylation is required for the establishment of front-to-back polarity and regulation of FAs and stress fibers

While our data demonstrate that Rap2 is required for regulation of lamellipodia dynamics, establishment of front-to-back polarity, and FA dynamics, it remains to be determined whether Rab40/CRL5-dependent mono-ubiquitylation of Rap2 is involved in regulating all these processes. To test whether Rap2 mono-ubiquitylation is required for these cell migration related functions, we generated MDA-MB-231 Rap2-KO lines rescued by overexpression of ether GFP-Rap2a-WT or a GFP-Rap2a-K3R mutant (K117R, K148R, K150R mutations that prevent ubiquitylation by the Rab40/CRL5 complex (Duncan et al., 2022)). Microscopy was then used to visualize the actin cytoskeleton and analyze the formation of front-to-back polarity. The loss of front-to-back polarity observed in Rap2-KO lines was mostly restored to control cell levels by expressing GFP-Rap2a-WT, while GFP-Rap2a-K3R expression failed to rescue any of these defects (Figs. 5A-C, 2A-C).

**Figure 5:**
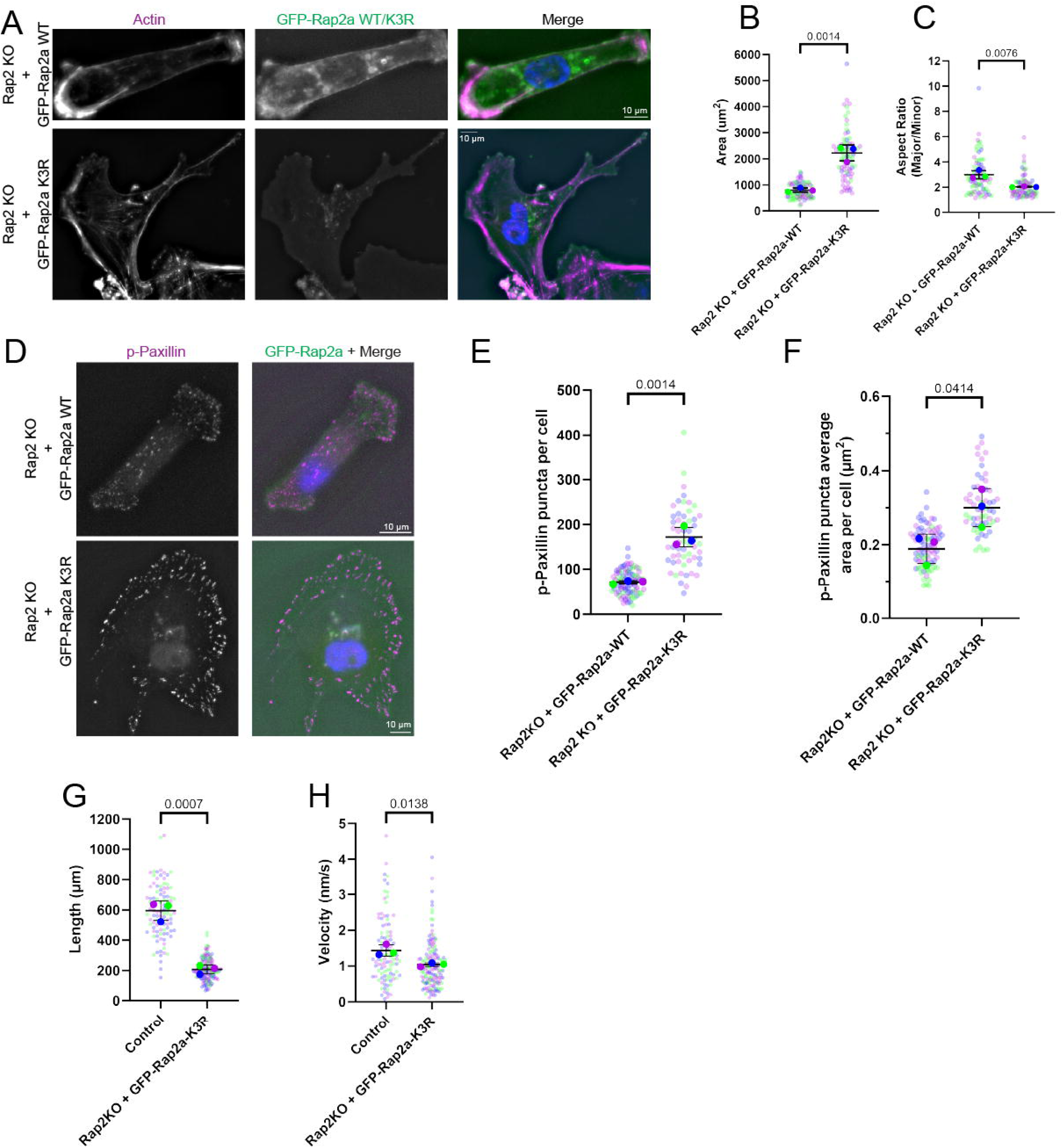
Mono-ubiquitylation is necessary for Rap2 function during cell migration. **(A-C)** Phalloidin-Alexa596 staining of Rap2KO cells rescued with either a GFP-Rap2a-WT or GFP-Rap2a-K3R construct. Quantification of cell size **(B)** and cell aspect ratio **(C). (D-F)** Control and Rap2-KO MDA-MB-231 cells were stained with phalloidin-Alexa596 (Actin) and an anti-phosphorylated(p)-Paxillin antibody. **(E-F)** Quantification of focal adhesions from cells shown in panel **(D)**. Panel **(E)** shows the average number of p-Paxillin puncta per cell and panel **(F)** shows the average size of p-Paxillin puncta per cell. **(G-H)** Quantification of tracks from random migration assays of Rap2KO cells rescued with either a GFP-Rap2a-WT or GFP-Rap2a-K3R construct. Quantifications show the total length of the track the cells migrated **(G)** and the velocity at which the cells migrated **(H)**.

Next, we sought to understand if Rap2 mono-ubiquitylation is also needed for the function of Rap2 in regulating FAs and actin stress fibers. To that end, we used an anti-phospho-paxillin antibody to analyze the size and number of FAs in GFP-Rap2a-WT or GFP-Rap2a-K3R expressing Rap2-KO cells. While GFP-Rap2a-WT overexpression rescued the Rap2 KO-induced FA defects, overexpression of GFP-Rap2a-K3R mutant failed to rescue any FA phenotypes (Fig. 5D-F, Fig. 4D-F).

Because GFP-Rap2a-K3R rescue lines were unable to rescue morphological or FA defects, we predicted that GFP-Rap2a-K3R rescue lines would also be unable to rescue Rap2 KO induced loss of migration. Accordingly, we repeated random migration assays with control and GFP-Rap2a-K3R rescue lines. GFP-Rap2a-K3R was not seen to rescue cell migration velocity and length (Fig. 5G-H) and mimicked migration characteristics of Rap2-KO cells (Fig. 1B-C). Thus, our data show that mono-ubiquitylation is necessary for Rap2 function during cell migration.

### Ubiquitylated Rap2 is enriched at the plasma membrane and lamellipodia

Because of the importance of Rap2 ubiquitylation in regulating cell migration, we next sought to visualize the subcellular localization of mono-ubiquitylated Rap2. Our previous work has shown that inhibition of Rap2 ubiquitylation causes its localization to shift from the plasma membrane and lamellipodia to lysosomes (Duncan et al., 2022), leading to predictions that mono-ubiquitylated Rap2 is likely present at the lamellipodia and plasma membrane. Consistent with this hypothesis, we have also shown that Rab40/CRL5 is present at the lamellipodia of migrating cells (Duncan et al., 2022; Linklater et al., 2021). However, these predictions fail to directly visualize the localization of ubiquitylated Rap2, leaving alternative explanations for the role of Rap2 ubiquitylation in trafficking and localization. To address this knowledge gap, we performed Proximity Ligation Assay (PLA) using anti-GFP and anti-ubiquitin antibodies in MDA-MB-231 cell lines overexpressing either GFP-Rap2a-WT or GFP-Rap2a-K3R. Consistent with our previous reports that Rab40/CRL5 mono-ubiquitylates Rap2 on K117, K148, and/or K150, cells expressing GFP-Rap2a-WT had more PLA puncta than cells expressing GFP-Rap2a-K3R (Fig. 6A-C). Importantly, cells assayed as controls using only anti-GFP or anti-ubiquitin antibodies (Fig. 6A-C, S2A-B) showed very little PLA signal (Fig. 6C, S2A). Furthermore, the greater number of PLA puncta in GFP-Rap2a-WT expressing cells was not a result of change in cell size (Fig. S2B). This suggests that mono-ubiquitylation of Rap2 by the Rab40/CRL5 ubiquitin ligase complex is the predominant form of Rap2 ubiquitylation in a cell.

**Figure 6:**
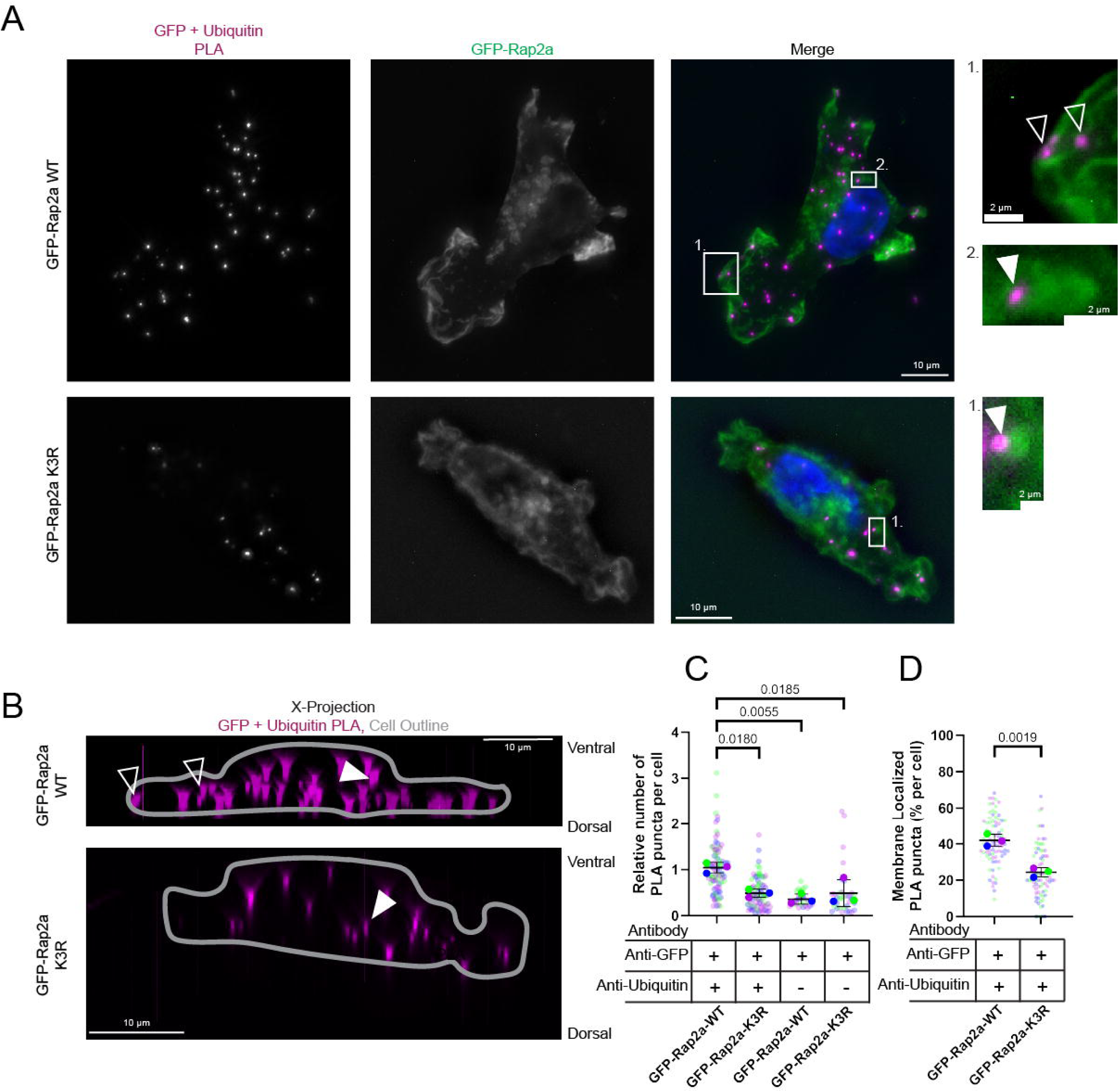
Rap2 is mono-ubiquitylated at the plasma membrane. **(A-D)** Proximity Ligation Assay (PLA) of MDA-MB-231 cells expressing either GFP-Rap2a-WT or GFP-Rap2a-K3R. In zoomed in sections, unfilled arrows highlight PLA puncta on the plasms membrane while filled arrows show PLA puncta on GFP-Rap2a vesicles **(A).** Panel **(B)** shows the same images as **(A)** but projected over the images X-axis. The GFP-Rap2a channel was traced to show the cell outline and displayed overtop the PLA channel. Arrows identify the same PLA puncta as shown in **(A).** Quantification of PLA puncta showing the average number of PLA puncta per cell **(C)** including PLA control data from **(Fig. S2),** all normalized per replicate to the median of the GFP-Rap2a-WT PLA data, and the localization of PLA puncta to the membrane **(D).**

Next, we asked where PLA puncta are enriched in cells expressing GFP-Rap2a-WT, as an indication of the localization of ubiquitylated Rap2. As shown in figure 6A-B and D, the majority of the PLA puncta were localized to the plasma membrane, especially in the areas of membrane ruffling, with some localization to GFP-positive vesicles. In contrast, the few PLA puncta observed in GFP-Rap2a-K3R expressing cells were localized in the cytoplasm (Fig. 6A-B and D, Fig. S2C). Thus, consistent with our previous reports that Rap2 is ubiquitylated by plasma membrane bound Rab40/CRL5 complex, Rap2 localized to the plasma membrane is mono-ubiquitylated and remains ubiquitylated through the internalization process.

### Ubiquitylation facilitates Rap2 retention at the lamellipodia plasma membrane

Our PLA experiment suggests that mono-ubiquitylated Rap2 is localized to the lamellipodia and plasma membrane and that the loss of ubiquitylation results in Rap2 accumulation in lysosomes (Fig 7A, (Duncan et al., 2022)). However, it remained unclear how ubiquitylation mediates Rap2 localization to the lamellipodia and plasma membrane.

**Figure 7:**
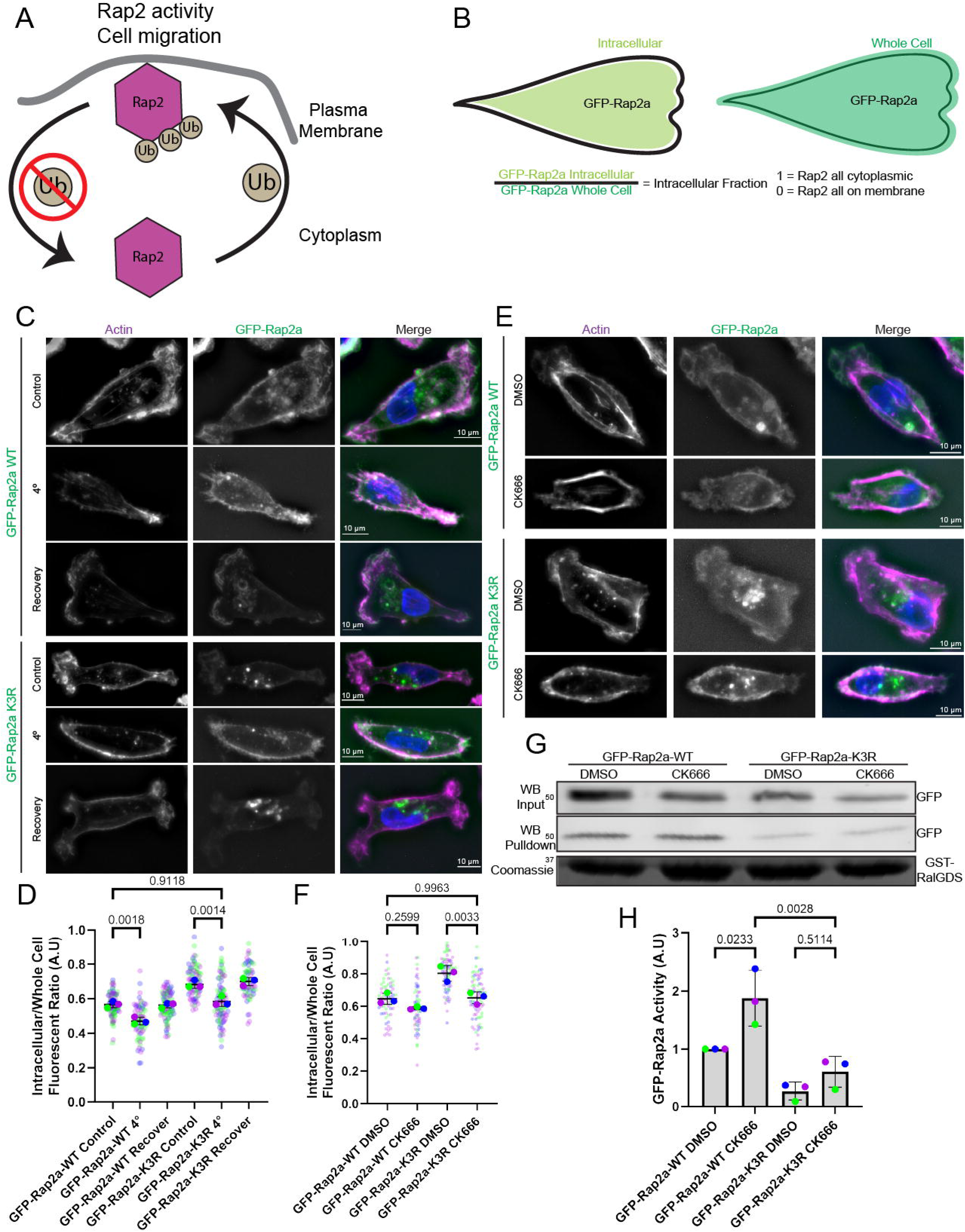
Mono-ubiquitylation facilitates Rap2 retention at the plasma membrane. **(A)** Schematic representation of Rap2 tri-mono-ubiquitination cycle. **(B)** Schematic representation showing the process used to calculate GFP-Rap2a plasma membrane localization. **(C-D)** MDA-MB-231 cells expressing either GFP-Rap2a-WT or GFP-Rap2a-K3R were subject to 4LC treatment to inhibit Rap2 internalization from the plasma membrane. 37LC temperature recovery was used to reinitiate Rap2 internalization via endocytosis/macropinocytosis. Quantification of Rap2 membrane localization is shown in **(D). (E-F)** MDA-MB-231 cells expressing either GFP-Rap2a-WT or GFP-Rap2a-K3R were treated with CK666 to inhibit macropinocytosis-dependent internalization of Rap2. Quantification of Rap2 membrane localization is shown in **(F). (G-H)** GST-RalGDS bead pulldown assays were used to test the levels of activated GFP-Rap2a-WT and GFP-Rap2a-K3R after CK666 treatment. Rap2 binding to GST-RalGDS was assessed by western blot and Coomassie blue staining was used to confirm the loading of GST-RalGDS into cell lysates. Quantification of western blots of GFP-Rap2a binding to GST-RalGDS normalized to GFP-Rap2a levels in the lysate **(H).**

To investigate this mechanism, we hypothesized two possible explanations for the role of ubiquitylation in Rap2 accumulation at the lamellipodia membrane. First, mono-ubiquitylation may mediate Rap2 trafficking to the plasma membrane. Second, mono-ubiquitylation may inhibit Rap2 removal from plasma membrane, thus, increasing its dwell-time at the lamellipodia.

Previous work has shown that Rap2 is predominately membrane-associated and is removed from the plasma membrane by macropinocytic vesicles that form at the actively ruffling lamellipodia (Duncan et al., 2022). Accordingly, to test these hypotheses, we used multiple techniques to inhibit Rap2 macropinocytic internalization from the plasma membrane and assessed the effect on GFP-Rap2a-WT and GFP-Rap2a-K3R localization at the plasma membrane (Fig. 7B). First, we inhibited endocytosis/macropinocytosis by incubating cells at 4[C. 4[C-treatment resulted in increased accumulation of both GFP-Rap2a-WT and GFP-Rap2a-K3R on the plasma membrane (Fig. 7C-D). Notably, moving cells back to 37[C led to a rapid removal of GFP-Rap2a-K3R from the plasma membrane and its accumulation in endosomes/lysosomes (Fig. 7C-D).

While effective in inhibiting endocytic/macropinocytic internalization, 4[C treatment has the potential to alter many other cellular functions such as enzymatic activity and protein interactions. To more directly inhibit macropinocytosis, we used CK666, the well-established inhibitor of the Arp2/3 complex. CK666 treatment inhibits branched actin polymerization, a necessary step in ruffling-dependent macropinocytosis (Mylvaganam et al., 2021). As shown in figure 7E-F, CK666 inhibition of macropinocytosis increased plasma membrane accumulation of both GFP-Rap2a-WT and GFP-Rap2a-K3R. Since inhibition GFP-Rap2a-K3R internalization rescues the Rap2a-K3R localization defect, mono-ubiquitylation of Rap2 does not play a role in Rap2 trafficking to the membrane. Instead, ubiquitylation likely functions to inhibit Rap2 internalization, increasing Rap2 dwell-time at the lamellipodia plasma membrane.

It is well-established that Rap2 GEFs are located at the plasma membrane (Bos et al., 2007; Kumar et al., 2018; Sartre et al., 2023). Thus, it was proposed that targeting Rap2 to the plasma membrane leads to its binding to Rap2 GEFs, resulting in Rap2 activation. We have previously shown that Rap2-K3R mutation blocks its activation (Duncan et al., 2022), raising an intriguing possibility that increasing Rap2-K3R dwell-time at the plasma membrane may rescue the Rap2-K3R activation defect. To test this hypothesis, we used CK666 to induce Rap2-K3R accumulation on the plasma membrane and tested Rap2 activity by performing Rap pulldown assays using GST-RalGDS (known Rap2 effector protein that binds Rap2-GTP) beads. GFP-Rap2a-WT was found to be more active when treated with CK666 (Fig. 7G-H), consistent with the idea that increased plasma membrane dwell-time increases Rap2 activation. However, CK666 treatment of cells overexpressing GFP-Rap2a-K3R only slightly increased its activation (Fig. 7G-H) indicating that forced membrane retention and rescue of GFP-Rap2a-K3R localization is not sufficient to rescue GFP-Rap2a-K3R activation. Thus, mono-ubiquitylation of Rap2 likely directly regulates GEF-dependent activation of Rap2.

### Mono-ubiquitylation regulates GEF-dependent Rap2 activation

Our data suggest that inhibition of Rab40/CRL5-dependent mono-ubiquitylation inhibits Rap2 activation (GTP binding) and decreases Rap2 dwell-time at the lamellipodia membrane.

Furthermore, we show that while we can rescue Rap2-K3R targeting to the plasma membrane by inhibiting macropinocytosis, it does not rescue defects in Rap2-K3R activation. These data indicate that ubiquitylation dependent GTP-loading of Rap2, likely through interaction with it cognitive GEF, is what maintains Rap2 localization at the lamellipodia membrane. To further test this hypothesis, we treated cells overexpressing GFP-Rap2a-WT or GFP-Rap2a-K3R with 8-bromo-cAMP, a cell permeable cAMP analog (Wang and Adjaye, 2011). cAMP stimulates activation of EPAC1, a known plasma membrane-associated GEF for Rap2 (Kumar et al., 2018; Sartre et al., 2023). As expected, we show that 8-bromo-cAMP treatment resulted in an increase of GFP-Rap2a-WT on the plasma membrane, consistent with the hypothesis that GTP-loading is what maintains Rap2 at the lamellipodia membrane (Fig. 8A-B). Importantly, 8-bromo-cAMP treatment did not change cytoplasmic localization of GFP-Rap2a-K3R (Fig. 8A-B), indicating that ubiquitylation is needed for Rap2 to be activated by EPAC1.

**Figure 8:**
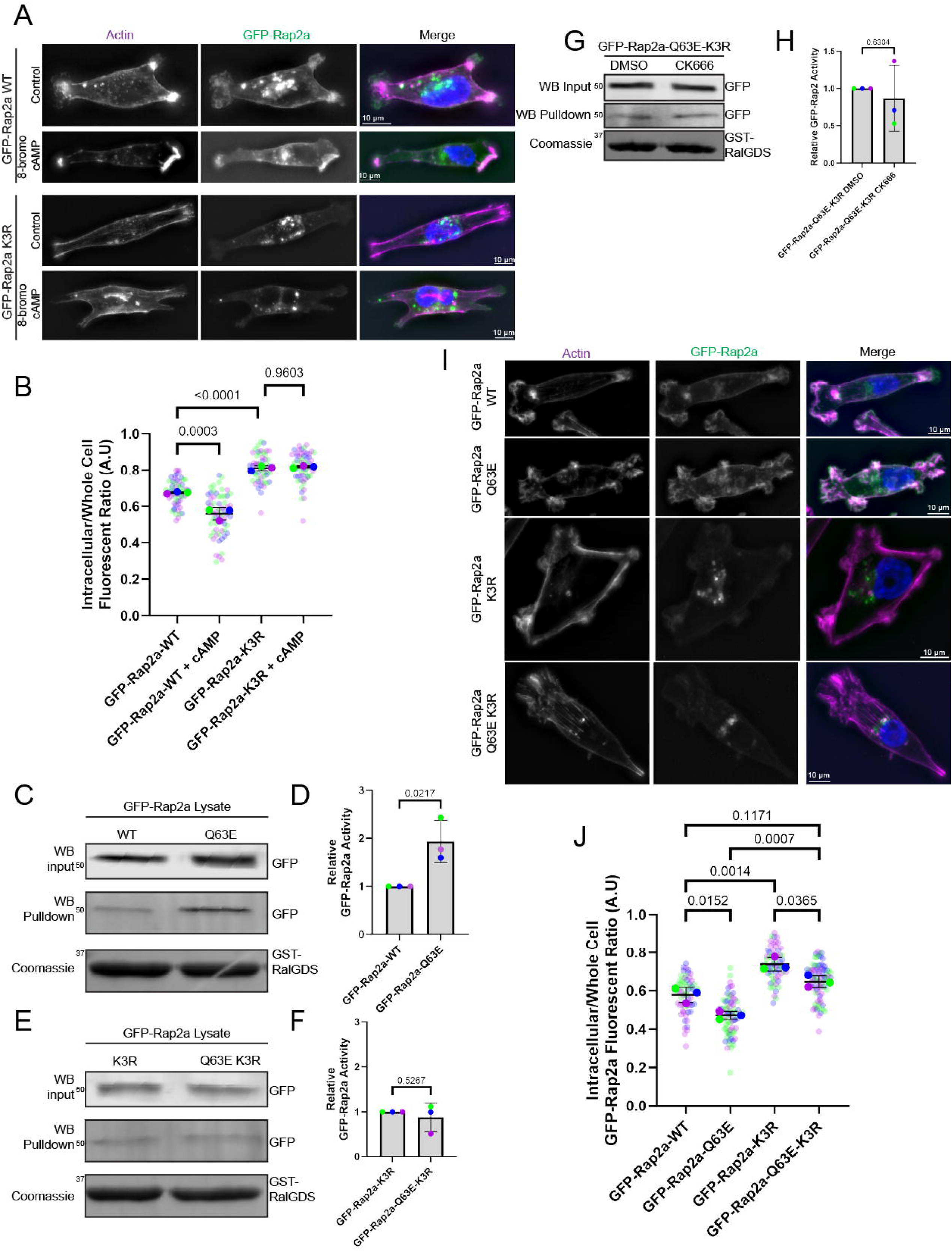
Mono-ubiquitylation mediates GEF-dependent activation of Rap2. **(A-B)** 8-bromo-cAMP was added to MDA-MB-231 cells expressing either GFP-Rap2a-WT or GFP-Rap2a-K3R in order to increase activity of a plasma membrane-associated Rap GEF and accordingly, increase Rap2 activity. Quantification of Rap2 membrane localization **(B). (C-F)** GST-RalGDS bead pulldown assays were used to measure the levels of activated GFP-Rap2a-WT and GFP-Rap2a-Q63E **(C)** and GFP-Rap2a-K3R and GFP-Rap2a-Q63E-K3R **(E)**. Quantification of GFP-Rap2a binding to GST-RalGDS beads was normalized to GFP-Rap2a levels in the lysate **(D,F). (G-H)** GST-RalGDS bead pulldowns were used to assess how CK666 affects GFP-Rap2a-Q63E-K3R activity. Quantification of GFP-Rap2a binding to GST-RalGDS beads was normalized to GFP-Rap2a levels in the lysate **(H). (I-J)** Localization of constitutively active GFP-Rap2a constructs in MDA-MB-231 cells. Quantification of Rap2 membrane localization **(J).**

We next decided to further analyze the role of Rap2 GTP-loading by generating the constitutively active Rap2 mutants, Rap2a-G12V and Rap2a-G12V-K3R (Janoueix-Lerosey et al., 1998; Lerosey et al., 1991). Unexpectedly, GFP-Rap2a-G12V localization phenocopied GFP-Rap2a-WT while GFP-Rap2a-G12V-K3R localization mimicked GFP-Rap2a-K3R localization (Fig. S3A-B). To assess the expected increase in activation of GFP-Rap2a-G12V, we performed active Rap2 pulldown activity assays using GFP-Rap2a-WT, GFP-Rap2a-G12V, and GFP-Rap2a-G12V-K3R cell lysates. While we could pull-down GFP-Rap2a-WT, both GFP-Rap2a-G12V and GFP-Rap2a-G12V-K3R failed to pull-down with GST-RalGDS beads (Fig. S3C), suggesting that the G12V mutation may not generate constitutively active Rap2 in our cells. The original study investigating the Rap2 G12V mutation was performed using purified protein, without the presence of Rap2 GAPs and GEFs. This work found that the Rap2 G12V mutation decreased innate Rap2 GTPase activity in the absence of GAPs (Lerosey et al., 1991). Thus, it is possible that in the cellular environment Rap2-G12V mutant does not function as a constitutively active Rap2. Consistent with this hypothesis, previous work identifying Rap2 binding partners showed that the Rap2-G12V mutant binds less efficiently to a Rap2 effector protein (Janoueix-Lerosey et al., 1998).

Since we could not use the Rap2-G12V mutant in our studies, we sought to create another constitutively active Rap2 mutation. Rap GTPase family member Rap1 has been shown to be constitutively activate upon a Q63E mutation (Dao et al., 2009; Zhang et al., 2014). Similar Q-to-E mutations were also shown to generate constitutively active mutants in other small monomeric GTPases, such as Ras and Rab GTPases (Gopal Krishnan et al., 2020; Langemeyer et al., 2014; Prior et al., 2012). Thus, we made the Q63E mutation in Rap2 to create a GFP-Rap2a-Q63E mutant and performed GST-RalGDS pulldown assays to test the activity of the Rap2-Q63E mutant. As shown in the figure 8C-D, GFP-Rap2a-Q63E bound GST-RalGDS beads stronger than GFP-Rap2a-WT, showing that Rap2-Q63E is an activating mutation. Accordingly, we tested the effect of tri-mono-ubiquitylation on activation of Rap2-Q63E by creating a GFP-Rap2a-Q63E-K3R mutant. Using GST-RalGDS pulldown assays we found that, like the GFP-Rap2a-K3R mutant, GFP-Rap2a-Q63E-K3R is largely inactive (Fig. 8E-F). It was proposed that Q mutations in the switch-II region of Ras GTPases block their interaction with GAPs (Prior et al., 2012; Vetter and Wittinghofer, 2001), thus locking them in the GTP-bound state. As such, the Rap2-Q63E mutant is likely unable to be deactivated by GAPs promoting its presence in the GTP-bound state. Since the combination of the Rap2-Q63E-K3R mutations render Rap2 inactive, our GST-RalGDS pulldown data then suggest that blocking Rap2 ubiquitylation inhibits Rap2 interaction with GEFs, preventing Rap2-Q63E-K3R activation.

To further test whether ubiquitylation is necessary for GEF dependent Rap2 activation, we treated cells expressing GFP-Rap2a-Q63E-K3R with CK666 to force their proximity with membrane localized GEFs and performed GST-RalGDS pulldowns. While decreasing internalization from the plasma membrane through CK666 treatment increases the activation of GFP-Rap2a-WT, likely as a result of increasing Rap2 localization with its GEFs (Fig. 7G-H),

CK666 treatment did not increase GFP-Rap2a-Q63E-K3R activation (Fig. 8G-H). Thus, even a Rap2-Q63E mutant first needs to be mono-ubiquitylated in order to be activated when at the lamellipodia membrane. As such, our data suggest that ubiquitylation of Rap2 is a prerequisite for GEF-dependent Rap2 activation.

Next, we sought to understand if ubiquitylation dependent Rap2 activation by GEFs also regulated Rap2 localization. As such, we observed the localization of our GFP-Rap2a-Q63E and GFP-Rap2a-Q63E-K3R constructs in MDA-MB-231 cells. GFP-Rap2a-Q63E was observed to localize more strongly to the lamellipodia and plasma membrane than GFP-Rap2a-WT (Fig. 8I-J), suggesting that increased activity as a result of decreased Rap2-GAP interaction increases Rap2 localization at the lamellipodia plasma membrane. Interestingly, the Q63E mutation did not rescue localization of the GFP-Rap2a-Q63E-K3R (Fig. 8I-J), further supporting the hypothesis that ubiquitylation dependent Rap2 activation is necessary for Rap2 maintenance at the lamellipodia plasma membrane.

Finally, using our Q63E mutants, we wanted to assess the role of increased Rap2 activity in the lamellipodia by analyzing GFP-Rap2a-Q63E dynamics in lamellipodia ruffles (Video 8). Consistent with the idea that an increase in Rap2 activity inhibits ruffle-associated RhoA, GFP-Rap2a-Q63E ruffle retraction in in the lamellipodia was slower and shorter as compared to GFP-Rap2a-WT ruffles (Fig. S4A-B). Furthermore, in GFP-Rap2a-Q63E expressing cells lamellipodia ruffling appears to be less periodic and more disorganized (Fig. S4C).

Altogether, our data suggest that Rap2 activation regulates its localization by increasing Rap2 dwell-time at the lamellipodia membrane. Accordingly, it is predicted that CK666 induced accumulation of wild type Rap2 on the membrane should be maintained after CK666 removal while the Rap2-K3R mutant should be removed from the membrane as it was never able to be activated. In accordance with these predictions, washout of CK666 results in rapid GFP-Rap2a-K3R removal from the plasma membrane while GFP-Rap2a-WT remained at the lamellipodia membrane (Fig. S5A-B).

## DISCUSSION

Cell migration is a vital process relying on the precise coordination of many intracellular molecular mechanisms to establish front-to-back polarity and create functional migratory structures. Here, we describe the mechanism by which Rab40/CRL5 dependent mono-ubiquitylation regulates the spatiotemporal dynamics of Rap2. We also demonstrate that Rap2 functions as a RhoA inhibitor during retraction of lamellipodia ruffles, thus promoting lamellipodia dynamics and cell migration.

### The Rap2 GTPase as a major facilitator of lamellipodia dynamics

Mesenchymal cell migration is hallmarked by the formation of a lamellipodia at the leading edge of the cell. Canonically, the lamellipodia defines the direction of cell migration and forms as a result of polarized Rac1 activation at the leading edge of the cell. Rac1 then activates the Arp2/3 complex which nucleates branched actin polymerization, creating the pushing force at the front of the lamellipodia membrane that drives membrane extension (Bisi et al., 2013; Schaks et al., 2019; Suraneni et al., 2012). However, recently it was shown that RhoA activity is also needed at the lamellipodia to facilitate ruffling and allow for lamellipodia dynamic extension and retraction cycles (Kurokawa and Matsuda, 2005; O’Connor and Chen, 2013; O’Connor et al., 2000). We show that mono-ubiquitylated Rap2 regulates lamellipodia dynamics by acting as a negative RhoA regulator. Specifically, we propose that Rap2 recruitment to retracting lamellipodia ruffles inhibits RhoA-induced acto-myosin contraction, terminating ruffle retraction, thus promoting branched actin polymerization and lamellipodia extension (Fig. 9A). Consistent with this hypothesis, Rap2 KO increases ruffle retraction distance and decreases ruffling periodicity. This decrease in lamellipodia ruffling likely contributes to the inhibition of cell migration in Rap2-KO cells. It remains unclear, however, how Rap2 mediates inactivation of RhoA. It is likely that Rap2 recruits RhoA GAPs to the retracting ruffles, thus leading to inactivation of RhoA. Consistent with this hypothesis, Rap2 has been implicated in regulation of RhoA GAP localization in other cellular contexts (Pannekoek et al., 2013; Post et al., 2015).

**Figure 9:**
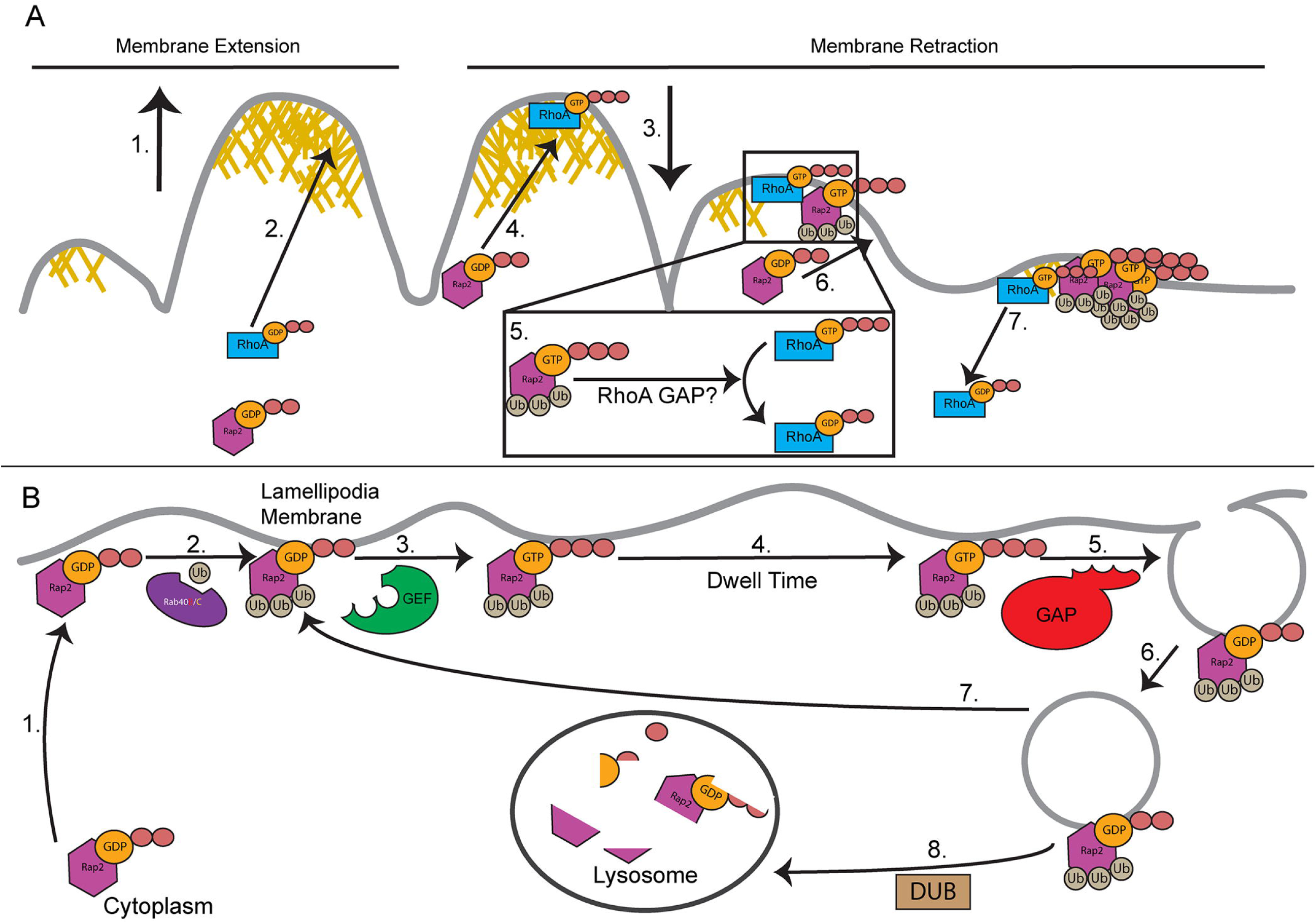
Mono-ubiquitylation mediates Rap2 activation, localization, and function during cell migration. **(A)** Proposed <u>model of Rap2 function during cell migration</u>. Membrane extension at the lamellipodia is hallmarked by branched actin polymerization, pushing the membrane outwards **(1).** At the end of the extension event, RhoA is recruited to the membrane and is activated **(2)** where it functions to mediate ruffle retraction **(3).** As the ruffle retracts, Rap2 is recruited to the retracting ruffle **(4).** Once localized, Rap2 is activated and inhibits RhoA activity, likely through recruitment of a RhoA GAP **(5).** Continued Rap2 recruitment **(6)** will inactivate RhoA, limiting the extent of ruffle retraction and removing RhoA activity from the membrane **(7).** Rap2 will then be deactivated and removed from the membrane allowing for another ruffle extension/retraction cycle. **(B)** Proposed <u>model of the role of mono-ubiquitination in regulating Rap2 localization and</u> <u>activation at lamellipodia membrane</u>. Rap2 is trafficked from the cytoplasm to the lamellipodia membrane **(1).** On the membrane, it is met and tri-mono-ubiquitinated by the Rab40/CRL5 E3 ubiquitin ligase complex **(2).** Once ubiquitinated, Rap2 is activated by a GEF, **(3).** Active (GTP-bound) Rap2 is retained at the plasma membrane where it functions to regulate lamellipodia dynamics **(4).** After Rap2 is inactivated by GAPs **(5)**, it is internalized through macropinocytosis **(6).** When internalized into an early endosomes Rap2 can either be recycled back to the plasma membrane were its re-activated **(8)** or trafficked to the lysosome for degradation **(9).** We speculate that the decision for Rap2 recycling or trafficking to the lysosome is based on whether Rap2 is deubiquitinated by a deubiquitinase (DUB) leading to lysosomal degradation or whether it maintains its ubiquitination and is recycled.

Notably, many RhoA GAPs are known to exist, multiple of which have been implicated in regulating cell migration (Mosaddeghzadeh and Ahmadian, 2021). Further studies will be needed to identify RhoA GAPs that bind to or is regulated by Rap2 and to understand the molecular machinery governing Rap2-RhoA GAP interactions during cell migration.

Our study also shows that the effect of Rap2 loss was not limited to defects in lamellipodia dynamics. Specifically, FAs were seen to be more stable and acto-myosin stress fibers were more prevalent/contractile in Rap2-KO cells. Importantly, the connection between FAs and stress fibers is well-established as stress fibers anchor to FAs. Furthermore, increased stress fiber contractility leads to activation and stabilization of FAs (Bachmann et al., 2022; Burridge and Guilluy, 2016; Wolfenson et al., 2011). Finally, regulation of FA assembly/disassembly dynamics is known to play a key role in regulating cell migration (Chen, 1979; Mishra and Manavathi, 2021; Wehrle-Haller, 2012). Both FAs and acto-myosin stress fibers are structures regulated by RhoA signaling (Hotulainen and Lappalainen, 2006; Julian and Olson, 2014). Thus, Rap2 regulated RhoA activity in lamellipodia is necessary for maintaining FAs/stress fibers. Defects in all these structures work together as strong driving factors to explain the loss of migration observed in Rap2-KO cells.

### Mono-ubiquitylation mediates Rap2 targeting to the lamellipodia membrane

Rab40/CRL5 E3 ubiquitin ligase-dependent mono-ubiquitylation of Rap2 has been shown to be necessary for its localization to the plasma membrane and lamellipodia ruffles (Duncan et al., 2022). However, it remained unknown where and when Rap2 is ubiquitylated and how this ubiquitylation mediates Rap2 targeting to the lamellipodia. In this study we used PLA to demonstrate that Rap2 is ubiquitylated at the plasma membrane and that mono-ubiquitylated Rap2 is specifically enriched at the lamellipodia ruffles. This observation is consistent with our previous study showing that Rab40/CRL5 complex is also enriched at the lamellipodia ruffles (Duncan et al., 2022; Linklater et al., 2021). In line with these results, we propose that, upon delivery to lamellipodia plasma membrane, Rap2 is mono-ubiquitylated by Rab40/CRL5 complex. This mono-ubiquitylation is required for Rap2 to accumulate and function at the lamellipodia. Interestingly, some ubiquitylated Rap2 can also be observed at endocytic vesicles. This observation suggests that in addition to controlling Rap2 localization to the membrane, ubiquitylation of Rap2 may also regulate Rap2 function beyond the plasma membrane and lamellipodia. Further work, however, will be necessary to conclusively answer these questions.

The observation that ubiquitylated Rap2 is enriched at the lamellipodia membrane raises the question of how mono-ubiquitylation mechanistically mediates Rap2 targeting to the plasma membrane. Canonically, ubiquitylation is a signal for plasma membrane-associated proteins, such as tyrosine kinase receptors, to be internalized and trafficked to the lysosome for degradation (Clague and Urbé, 2017; Critchley et al., 2018; Tanner et al., 2019; Weissman et al., 2011). However, mono-ubiquitylation has also been shown to regulate membrane protein trafficking without promoting degradation (Choi et al., 2018; Date et al., 2022; Xu et al., 2017).

While mono-ubiquitination does not lead to Rap2 degradation, these past works supported the idea that ubiquitylation can be directly involved in regulating protein delivery to or removal from the plasma membrane. Our data suggest that ubiquitylation does not directly regulate Rap2 trafficking to the plasma membrane as defects in Rap2-K3R localization can be rescued by inhibiting Arp2/3-dependent lamellipodia ruffling that drives macropinocytosis-dependent internalization of Rap2 (Duncan et al., 2022). Furthermore, re-initiation of lamellipodia ruffling by washing out Arp2/3 inhibitor leads to rapid removal of Rap2-K3R from plasma membrane and its accumulation in endosomes and lysosomes. These results suggest that mono-ubiquitylation increases Rap2 dwell-time at the lamellipodia membrane by inhibiting its removal from the plasma membrane. However, more work is needed to understand the molecular mechanism by which mono-ubiquitylation might govern Rap2 internalization. Furthermore, how Rap2 is de-ubiquitylated and the resulting consequence of this de-ubiquitylation on Rap2 trafficking still needs to be determined.

### Mono-ubiquitylation is necessary for GEF-dependent Rap2 activation

Small protein GTPases are canonically thought of being activated by GEFs. For Ras and Rap GTPases, activation is driven by plasma membrane-associated GEFs. Active (GTP-bound) Rap GTPases are then maintained at the plasma membrane where they function to mediate signaling and localized cytoskeleton dynamics. Eventual inactivation by GAPs results in internalization of GTPases and their removal from the plasma membrane (Bos et al., 2007; Segev, 2011). Consistent with this idea, we showed that the overactivation of a Rap2 GEF increases Rap2 plasma membrane localization in a mono-ubiquitylation-dependent manner. We further showed that enhanced localization to the plasma membrane increases Rap2 activity, likely due to increased co-localization with its membrane localized GEFs. In contrast to Rap2-WT, increasing Rap2-K3R localization at the lamellipodia membrane only slightly increased its activation. Furthermore, the activation of EPAC1 with cAMP, a known plasma membrane-associated GEF for Rap2, did not rescue the activation defects in Rap2-K3R mutants. Thus, this suggests that mono-ubiquitylation of Rap2 may directly modulate its ability to bind and be activated by its cognitive GEF.

To further explore the requirement for ubiquitylation in Rap2 activation, we generated Rap2-Q63E and Rap2-Q63E-K3R, constitutively active Rap2 mutants. The Q63E mutation was shown to be constitutively active in Rap1 (Dao et al., 2009; Zhang et al., 2014) and equivalent mutations are constitutively active in other small monomeric GTPases (Gopal Krishnan et al., 2020; Langemeyer et al., 2014; Prior et al., 2012; Vetter and Wittinghofer, 2001). In Ras and Rap GTPases, the conserved Q residues is located in the switch II region and line the entrance to the GTP binding pocket where a GAP needs to insert its arginine finger to facilitate the GTP-to-GDP conversion reaction. Consequently, mutation of these Q residues hinders Ras and Rap interactions with their cognitive GAPs (Prior et al., 2012; Vetter and Wittinghofer, 2001).

Consistent with this, we show that the Rap2-Q63E mutation leads to its activation and enhanced targeting to the plasma membrane. Interestingly, the Rap2-Q63E-K3R mutant failed to localize to the plasma membrane. Furthermore, the levels of active Rap2-Q63E-K3R were found to be similar to Rap2-K3R, suggesting that mono-ubiquitylation is necessary for Rap2 activation, potentially through directly regulating Rap2 interactions with its GEF.

Based on these results, we propose a Rab40/CRL5-dependent mechanism that regulates Rap2 activity during cell migration (Fig. 9A). Once Rap2 is delivered to the lamellipodia membrane (Duncan et al., 2022), it is mono-ubiquitylated by a Rab40/CRL5 complex that is enriched at the lamellipodia (Duncan et al., 2022; Linklater et al., 2021). This ubiquitylation mediates Rap2 interaction and activation by its cognitive GEF. Active Rap2 is then retained at the plasma membrane where it functions to regulate lamellipodia dynamics during cell migration (Fig. 9B). Notably, Ras GTPase family member K-Ras was also shown to be mono-ubiquitylated at K147. It was also demonstrated that mono-ubiquitylation of K-Ras at K147 inhibits its interaction with its GAP, thus increasing K-Ras activation (Choi et al., 2018; Sasaki et al., 2011). While we show that Rap2 ubiquitylation functions to allow for GEF-dependent activation, it is intriguing that the Ras family GTPases seems to have evolved a mono-ubiquitylation dependent mechanism to regulate their activity.

This conserved mechanism presents an exciting area of future cell migration research. As major regulators of cell migration and actin dynamics, understanding how ubiquitylation facilitates Ras/Rap localization and activation will lead to new insights into fine tuning cell migration as well as providing new modifications to target for treating cell migration related disorders and diseases.

## MATERIALS AND METHODS

### Cell culture

MDA-MB-231 cells were cultured in 231 media (DMEM with 4.5 g/liter glucose, 5.84 g/liter L-glutamine, 1% sodium pyruvate, 1% nonessential amino acids, 1 µg/ml insulin, 1% penicillin/streptomycin, and 10% FBS). HEK293T cells were cultured in 293T media (DMEM with 4.5 g/liter glucose, 5.84 g/liter L-glutamine, 1% penicillin/streptomycin, and 10% FBS). All MDA-MB-231 stable cells lines used in this study were generated using lentivirus plasmid pLVX as described previously (Duncan et al., 2022). The Rap2-KO line had been previously created and validated (Duncan et al., 2022). Cell lines were routinely tested for mycoplasma. Additionally, all cell lines were authenticated in accordance with ATCC standards.

### Generation of lentiviral stable cell lines

Calcium Phosphatase was used to transfect HEK293T cells (50% confluent) with pLVX plasmid containing GFP-tagged gene of interest. After 6 hours the media was replaced with fresh 293T media. Cells were left for 48 hours to allow for the virus to accumulate in the media. Viral 293T media was collected, filtered through a 0.45 µm PVDF low binding syringe filter, and treated with polybrene (100 µg per 1 mL media). Viral 293T media was added to target MDA-MB-231 cells (50% confluent) for 2 hours. Media was replaced with 231 media and target cells were allowed to recover for 24 hours, then selected with puromycin (5 µg/mL). Expression for the GFP-tagged gene of interest in cell lines were then validated using Western Blotting.

### Transient transfection of MDA-MB-231 cells

Transient transfections were carried out using the X-tremeGENE transfection reagent following the manufacturer’s recommended protocol. For transfection with GFP-Paxillin, cells were seeded in 35 mm glass-bottom dishes and transfected the same day. Time-lapse imaging was performed the following day. For the transfection with the RhoA biosensor (dTom-2xrGBD), cells were plated in a 35mm glass-bottom dish like before, except transfection was performed the next day. Time lapse imaging was performed the day after transfection.

### Random migration assay

A single-cell migration assay was performed using live-cell imaging. Imaging was conducted on an OLYMPUS IX83 inverted confocal microscope with a temperature-controlled stage top, which was maintained at 37°C with 5% CO_2_. Cells were plated uniformly in a 35 mm glass-bottom Petri dish that was pre-coated with Fibronectin. Fibronectin coating was performed by allowing the Fibronectin solution to dry for 1 hour under UV light. Cells were then seeded and allowed to adhere overnight. The next day, the old culture medium was replaced with a fresh medium the next day before starting the live imaging experiment. The time-lapse imaging was automatically taken and set to take at 20-minute intervals, capturing a total of 36 frames, resulting in a 12-hour time-lapse movie.

### Stroboscopic Analysis of Cell Dynamics (SACED)

SACED was performed as previously described (Hinz et al., 1999). Cells were plated in collagen coated glass bottom dishes and allowed to adhere overnight. 1 hour before imaging, 231 media was buffered with 40mM HEPES. Imaging was performed using a 64X DIC objective. Control cells were selected for imaging if they exhibit distinct ruffling lamellipodia and defined polarity.

Due to the loss of polarity and lamellipodia in Rap2-KO cells, these cells were selected based off the presence of a ruffling, lamellipodia-like structure. Before image acquisition, the field of view was cropped and focused to the ruffling edges of the cells. Images were taken for 8 mins, at 1 frame a second, with an exposure time of 100 ms. This was repeated across three replicates with multiple cells per replicate (Fig. 1E-G). Only cells which maintained focus of the ruffling edge throughout the whole 8 min video were selected for analysis.

### Immunofluorescence staining

MDA-MB-231 cells were seeded onto 1X collagen-coated glass cover glass slips and grown in full growth media for approximately 24 hours. Cells were later washed with room temperature PBS (TBS in case of phospho-antibody staining) and fixed it with 4% Paraformaldehyde for 15 minutes at RT. Cells were then quenched for 5 minutes with Quench buffer (375 mg of glycine diluted in PBS), and incubated in Incubation buffer (1ml of FBS, PBS, 1% BSA, 200mg saponin) for 30 minutes. Primary antibodies were diluted with 1:100 or 1:50 with the incubation buffer and placed on cells 1 hour in a hydrated humidified chamber. Cells were washed twice, then the secondary antibody was diluted at 1:100 and incubated with cells for 30 minutes in at humidified chamber. Cells were treated with 1:1600-1:2000 Hoechst stain for 5 minutes, then washed twice with PBS. Coverslips were then mounted onto glass slides using Vectashield.

### Active RhoA pulldown assays

GST-RBD beads were generated as previously described (Guilluy et al., 2011) with the exception that a French Pressure Cell was used to lyse cells instead of a sonicator. For the activity assay, nearly confluent monolayers (80-90%) in 10 cm plates were washed with cold PBS containing 1mM MgCl_2_. After washing, cells were scraped in 500 µL of lysis buffer containing 50 mM Tris-HCl pH 7.6, 500 mM NaCl, 1% Tritonx100, 0.1% SDS, 0.5% sodium deoxycholate, and 10 mM MgCl_2_ (added right before use). Cells were lysed for 5 minutes on ice then clarified in a 4LC centrifuge at 14,800xg for 5 mins. 10 µL of the lysate was used as a protein input control and approximately 400 µL of lysate was added to 50 µg of GST-RBD beads. The mixture was rotated at 4LC for 30 minutes. Beads were washed 3x with 1 mL of cold wash buffer containing 50 mM Tris-HCl pH 7.6, 150 mM NaCl, 1% Tritonx100, and 10 mM MgCl_2_ (added right before use). All wash buffer was removed using a fine pipette tip and protein was eluted from beads in 35 µL of 1x SDS sample loading dye. Samples were separated by SDS-PAGE and analyzed by Coomassie staining and western blot. 20 µL of elution and 10 µL protein input controls were used for western blotting. 10 µL of elution was used for Coomassie staining to confirm the presence of GST-RBD.

### Cell lysis and Western blotting

Unless otherwise stated, cells were lysed on ice in buffer containing 20 mM Hepes, pH 7.4, 150 mM NaCl, 1% Triton X-100, and 1 mM PMSF. After 30 min, lysates were clarified at 15,000 *g* in a prechilled microcentrifuge. Supernatants were collected and analyzed via Bradford assay.

Lysate samples were prepared in 5X-SDS loading dye, boiled for 5 min at 95°C, and separated via SDS-PAGE. Gels were transferred onto 0.45-µm polyvinylidene difluoride membrane, followed by blocking for 1 hour in Intercept Blocking Buffer diluted in TBST 1:3. Primary antibodies (made in diluted Intercept Blocking Buffer) were incubated overnight at 4°C. Blots were then washed in TBST (Tris-buffered saline + 0.05% TWEEN) followed by incubation with IRDye fluorescent secondary antibody (diluted Intercept Blocking Buffer) for 1 hour at room temperature. Blots were washed once again with TBST before final imaging on a Li-Cor Odyssey CLx.

### Proximal Ligation Assay (PLA)

MDA-MB-231 cells stably expressing GFP-Rap2a wt or GFP-Rap2a K3R were plated onto collagen coverslips and left to adhere overnight. Cells were fixed in 10% PFA for 10 mins, quenched for 5 mins (375 mg glycine in 50 mL PBS), then permeabilized with 0.1% Triton X-100 for 5 mins. Following permeabilization, the Duolink proximal ligation assay (PLA) kit was used as described in the manufacturers protocol (Sigma Aldrich). Mouse anti-ubiquitin and rabbit anti-GFP primary antibodies were used together (or individually for controls). Experimental and control (GFP or ubiquitin only antibodies) cells were selected through visualization of GFP-Rap2a. Cells exhibiting front-to-back polarity with standard GFP-Rap2a fluorescence were selected for imaging. Three replicates were performed, each consisting of at least 20 cells for experimental conditions (both antibodies) or less for control conditions which contained minimal PLA puncta.

### 4°C and CK666 treatment

#### 4°C incubation

MDA-MB-231 cells stably expressing GFP-Rap2a-WT or GFP-Rap2a K3R were seeded on collagen coated coverslips. 24 hours later, 231 media was buffered with 20 mM Hepes, pH 7.4 and cells were divided into 3 groups: Control, 4L-incubation, and recovery. Control cells were placed in a standard incubator for 60 mins before being fixed with 4% paraformaldehyde. 4L-incubation cells were placed in a 4L cold room for 60 mins before fixing with 4% paraformaldehyde. Finally, recovery cells were placed in a 4L cold room for 60 mins, then placed in a standard incubator for 45 mins before fixing with 4% paraformaldehyde.

#### CK666 treatment

MDA-MB-231 cells stably expressing GFP-Rap2a-WT or GFP-Rap2a K3R were seeded on collagen coated coverslips. After 24 hours, cells were treated with 200 µM ck666 or DMSO (equal volume as ck666) as a control for 1 hour. Cells were then fixed with 4% paraformaldehyde and processed for immunofluorescence analysis. For CK666 washout experiments, after 1 hour of CK666 treatments, cells were carefully washed 3x with warmed PBS and placed with fresh media in the incubator for 20 mins to recover. Cells were then fixed with 4% paraformaldehyde.

For both experiments, cells with observable polarity were randomly selected for imaging through visualization of the actin cytoskeleton. Three biological replicates were performed with each replicate consisting of at least 20 randomly chosen cells (technical replicates).

### Active Rap2 pulldown assays

GST-RalGDS was purified as previously described (Duncan et al., 2022). MDA-MB-231 cells stably expressing GFP-Rap were grown in 10cm plates to 90% confluency, then harvested and pelleted (frozen if necessary). Pellets were lysed in buffer containing 20 mM Hepes, pH 7.4, 150 mM NaCl, 1% Triton X-100, 1 mM PMSF, and 5 mM iodoacetamide (DUB inhibitor) on ice for 30 mins. Lysate was clarified in chilled microcentrifuge at 21,000xg. Lysate was brought to equal concentration and volume in buffer containing 20mM Hepes pH 7.4 and 150mM NaCl. 20 µg of either GST (control) or GST-RalGDS was added to the lysate. Lysate controls (equal µg across conditions) were also taken for later use. Tubes with the GST proteins and lysate mixture were rotated for 60 mins at room temperature. 45 µL of glutathione beads (50% in PBS) were added to the tubes, and rotation was continued for 30 more minutes. Beads were then washed 5x in 1 mL buffer containing 20 mM Hepes, pH 7.4, 300 mM NaCl, and 0.1% Triton X-100. Protein was eluted from beads in 35 µL of 1x SDS sample loading dye, separated by SDS-PAGE and analyzed by Coomassie staining and western blot. 20 µL of elution and lysate controls were used for western blotting. 10 µL of elution was used for Coomassie staining to confirm the presence of GST/GST-RalGDS.

For assays with GTP locking of GFP-Rap protein (Fig S3C,D), the below steps were followed at room temperature prior to GST/GST-RalGDS addition. 1) 5mM EDTA was added to lysate for 5 mins. 2) 5mM GppCp was added to lysate for 5 mins. 3) 15 mM MgCl_2_ was added to lysate from a stock of MgCl_2_ diluted to 150 mM in water, pH 7.0 for 5 mins.

Western blots were analyzed using the gel analyzer tool in Fiji. Area from the pulldown was normalized to area of the lysate control to achieve a relative density measurement. Coomassie staining was not used for qualitative analysis.

### CK666 treatment for active Rap2 pulldown assays

The standard Rap activity pulldown assay was altered to minimize potential GAP activity. Cells expressing GFP-Rap2a were grown in 10 cm plates. Near confluent monolayers were treated with 200 µM CK666 or an equivalent volume of DMSO for 1 hour. During this time, GST-RalGDS was prebound to 45 µL of 50% GST bead slurry in 100 µL of buffer containing 20mM Hepes pH 7.4 and 150mM NaCl. Beads and protein were rotated at room temperature for 45 minutes after which the beads were pelleted and the supernatant was removed. Prebound beads were kept on ice.

After CK666 or DMSO treatment, cells monolayers were washed with buffer containing 20mM Hepes pH 7.4 and 150mM NaCl. Cells were then scraped into tubes with approximately 150 µL of lysis buffer containing 20 mM Hepes, pH 7.4, 150 mM NaCl, 1% Triton X-100, 1 mM PMSF, and 5 mM iodoacetamide (DUB inhibitor), Tubes were placed on ice for 5 minutes.

Lysate was clarified in chilled microcentrifuge at 21,000xg for 5 minutes. 400 µg of lysate was added to prebound GST-RalGDS beads and brought to an equal volume of 200 µL with buffer containing 20 mM Hepes, pH 7.4, 150 mM NaCl, 1 mM PMSF, and 5 mM iodoacetamide. Tubes were rotated at room temperature for 40 minutes.

After rotating, beads were then washed 3x in 1 mL buffer containing 20 mM Hepes, pH 7.4, 300 mM NaCl, and 0.1% Triton X-100. Protein was eluted from beads in 35 µL of 1x SDS sample loading dye, separated by SDS-PAGE and analyzed by Coomassie staining and western blot. 20 µL of elution and lysate controls were used for western blotting. 10 µL of elution was used for Coomassie staining to confirm the presence of GST/GST-RalGDS.

### cAMP treatment for immunofluorescence analysis

cAMP treatment of cells was used to test how activation and ubiquitylation of Rap2 interconnect. Cells stably expressing GFP-Rap2a wt or GFP-Rap2a K3R were seeded on collagen coated coverslips. After 24 hours, cells were treated with either 500 µM cAMP or water (equal volume as cAMP) as a control for 1 hour. Cells were then taken for immunofluorescence. Polarized cells were randomly selected for imaging through visualization of the actin cytoskeleton. Three replicates were performed with each replicated including at least 20 cells.

### Image acquisition

All fixed cell imaging was performed on a widefield inverted Zeiss Axiovert 200M microscope using a 63X oil-objective, QE charge-coupled device camera (Sensicam), and Slidebook v. 6.0 software (Intelligent Imaging Innovations). Images were taken as z-stacks with 0.5 µm step intervals. Where indicated images were deconvolved (Nearest Neighbors) using the Intelligent Imaging Innovations software. SACED and GFP-paxillin time-lapse images were acquired on the same Zeiss widefield with a 63X oil objective and a temperature-controlled stage. Further image processing was performed in Fiji software.

All random migration assays were performed using OLYMPUS IX83 inverted confocal microscope, with 4X air objective, equipped with a (HAMAMATSU) Camera controller.

Time-lapse imaging of GFP-Rap2a dynamics were collected on a Nikon Ti2-E inverted microscope equipped with 1.45 NA 100x CFI Plan Apo objective, Nikon motorized stage, Prior NanoScan SP 600 µm nano-positioning piezo sample scanner, CSU-W1 T1 Super-Resolution spinning disk confocal, SoRa disk with 1x/2.8x/4x mag changers, 488 nm, 560 nm, and 647nm laser, and a prime 95B back illuminated sCMOS camera.

### Image analysis

#### Random single-cell migration analysis

(Fig. 1A-C, 5G-H) was performed using the Manual Tracking Excellence-Pro software. Cells were tracked by their geographic center. Only cells that remind in focus throughout the whole time-lapse series and did not divide were tracked. Data was generated, and parameters like speed, length, and distance were acquired.

#### SACED analysis

(Fig. 1D-G, S1B-D) was performed by creating a kymograph in Fiji. Briefly, two lines were drawn through the ruffling section of the membrane at perpendicular angles to the membrane ruffles. The exact location of the line within the ruffling section was decided randomly. Two kymographs were created using the reslice tool; one for each line. For each kymograph, measurements were taken as previously described (Hinz et al., 1999) and as defined in Fig. S1A. The measurements from each event (ruffle retraction, membrane extension/retraction) were averaged together to produce one measurement for each kymograph. The average from the kymographs were then averaged together to create an average of each measurement per cell.

#### GFP-Rap2a dynamics

(Fig. 1J-K, 2A-C, S4B-C) were measured by creating kymographs in Fiji. Briefly, two lines were drawn through the ruffling section of the membrane at perpendicular angles to the membrane ruffles. The exact location of the line within the ruffling section was decided randomly. Two kymographs were created using the reslice tool; one for each line. Enrichment from the start to end of the ruffle (Fig. 1J-K) was measured from 3 ruffles per kymograph. The ruffles selected were the first complete ruffle visible, the last complete ruffle visible, and the ruffle most in the center of the kymograph. For each selected ruffle, a ROI was drawn around approximately the first third and last third of the ruffle. Mean grey value was measured from each ROI and the background mean grey value was subtracted from each measurement. The average mean grey value was found for each cell. Enrichment was defined as the 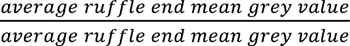. Ruffle retraction dynamics (Fig. S4B-C) were measured from the GFP-Rap2a using the same measurements as the SACED dynamics. When comparing GFP-Rap2a localization to the RhoA biosensor (dTom-2xrGBD, Fig. 2A-C), kymographs were smoothened two times. A line was drawn through the retracting ruffle of the GFP-Rap2a channel of the kymograph and the fluorescent intensity along the line was measured in both the GFP-Rap2a and RhoA biosensor channels. This was repeated for 3 ruffles from each cell measured (8 total cells).

#### Cell morphology analysis

(Fig. 3B-C, 5B-C) was performed using Fiji. Images were max projected to include all stacks in which the actin cytoskeleton was in focus. The actin cytoskeleton from the resulting projection was traced using the polygon selection tool to create an ROI of the cell perimeter. The area, fitted ellipse, and perimeter of the ROI was then measured. Area was defined as the area (µm^2^), circularity as 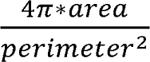, and aspect ratio as the 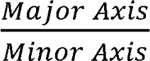 of the fitted ellipse of the ROI.

#### Cortactin enrichment analysis

(Fig. 3F-H) was performed using Fiji. Images were max projected to include all stacks in which the actin cytoskeleton was in focus. Cells with cortactin enrichment were defined as those with visible enrichment of cortactin fluorescence on a segment of the cell perimeter. Cortactin segments were measured in size by defining the percentage of the cell perimeter that they occupied. The actin cytoskeleton of the cell was traced using the polygon tool to create an ROI. The perimeter of the ROI was calculated. The segmented line tool was used to measure the length of the cortactin enriched segment. The segments percent of the perimeter was defined as 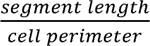. Cortactin enrichment was measured by drawing ROIs around the cortactin enriched region, a portion of the actin cortex (as defined by the actin channel), and the background fluorescence and mean grey value of the cortactin channel was measured and used for calculations. Enrichment was defined as 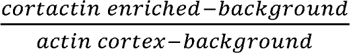. Some cells exhibited multiple cortactin enriched segments. The percentage of the perimeter and cortactin enrichment was calculated separately for each segment. Each measurement was used when calculating the average of a replicate.

#### Focal adhesion dynamics

(Fig 4A-D) The TrackMate plugin in Fiji was used to manually track GFP positive puncta (FAs). Puncta were tracked across each frame until they disappeared. Data for each puncta was generated by TrackMate

#### Focal adhesion count and size

(Fig. 4F-G, 5E-F) was measured in Fiji. 1-3 stacks where the p-paxillin signal (focal adhesions) was in focus were max projected. Images were cropped to only include one cell at a time and further analysis was performed blinded. The p-paxillin channel was threshold to apply a mask and create puncta for all focal adhesions in a cell. The analyze particle tool was then used to measure the size and number of each puncta was measured. Data was filtered to only include puncta larger than 0.04 µm^2^ and less than 2 µm^2^.

Filtered puncta size was averaged per cell. Puncta number and average size per cell was averaged across all cells to create the mean replicate value.

#### MyoIIb analysis

(Fig. 4H) was performed using line scans in Fiji. 1-2 images from z-stacks, where the myoIIb channel was in focus, were max projected. In the actin channel, a stress fiber was traced from start to end using the segmented line tool. The line was then transferred to the MyoIIb where the plot profile tool was used to take fluorescent intensity measurements from across the line. Fluorescent intensity of MyoIIb signal was plotted against its location along the stress fiber.

#### PLA analysis

(Fig. 6A-D, Fig. S2A-C) was performed using Fiji. All z-stack images were max projected to include all PLA puncta except puncta on the cell’s dorsal surface. The images were then cropped to only show one cell at a time. The polygon tool was used to trace both the outside of the GFP-Rap2a signal, creating a whole cell ROI, and the inside of the GFP-Rap2a signal, creating an intracellular ROI. A threshold was applied to the PLA channel to make masks of PLA puncta. Thresholding was done to exclude any residual signal from dorsal surface PLA puncta. The analyze particle tool was used to count the number of puncta greater than 0.04 µm^2^ in the whole cell ROI. To count membrane localized puncta, the intracellular ROI was then filled and the number of puncta remaining was counted in the same way. The percent of membrane localized PLA puncta was defined as 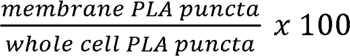. Vesicle associated puncta were defined using z-stack images. Vesicles were identified by eye and PLA puncta that were in focus in the same plane +/-1 were considered vesicle associated. Percent was calculated as 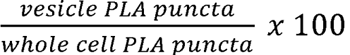.

#### *Intracellular/Whole cell fluorescence* ratio

(Fig. 7D,F, 8B,H, S3B, S5B) was calculated in Fiji using images that were max projected on the stacks where GFP-Rap2a signal was in focus. Per cell, the actin channel was used to define the cell perimeter. An ROI was traced on the outside of the actin channel and defined as the whole cell while another ROI was traced on the inside of the actin channel and defined as the intracellular ROI. The ROIs were then applied to the GFP-Rap2a channel and the integrated density was measured. ROIs were also used to measure the integrated density of the background fluorescence. Data was expressed as a ratio of integrated density which was defined as 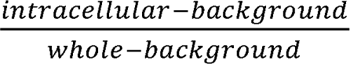 where a value of 1 would indicate all the signal was intracellular.

### Statistical analysis

Graphs are displayed as biological replicates (opaque dots) overlayed on technical replicates (translucent dots). Error bars represent the standard deviation of the biological replicates.

Replicates are displayed by color. Statistical analysis was performed on biological replicates which were calculated from the mean of the technical replicates. For all analyses comparing 3 or more conditions, a one-way ANOVA was used with comparisons run for each column against the mean of each other column. p-values of interest are displayed on graphs. For all analyses comparing 2 conditions, a student’s t-test was used with all significant p-values displayed. A paired t-test was used for Figure 1J and unpaired t-tests were used for all other t-tests.

**Table 1:**
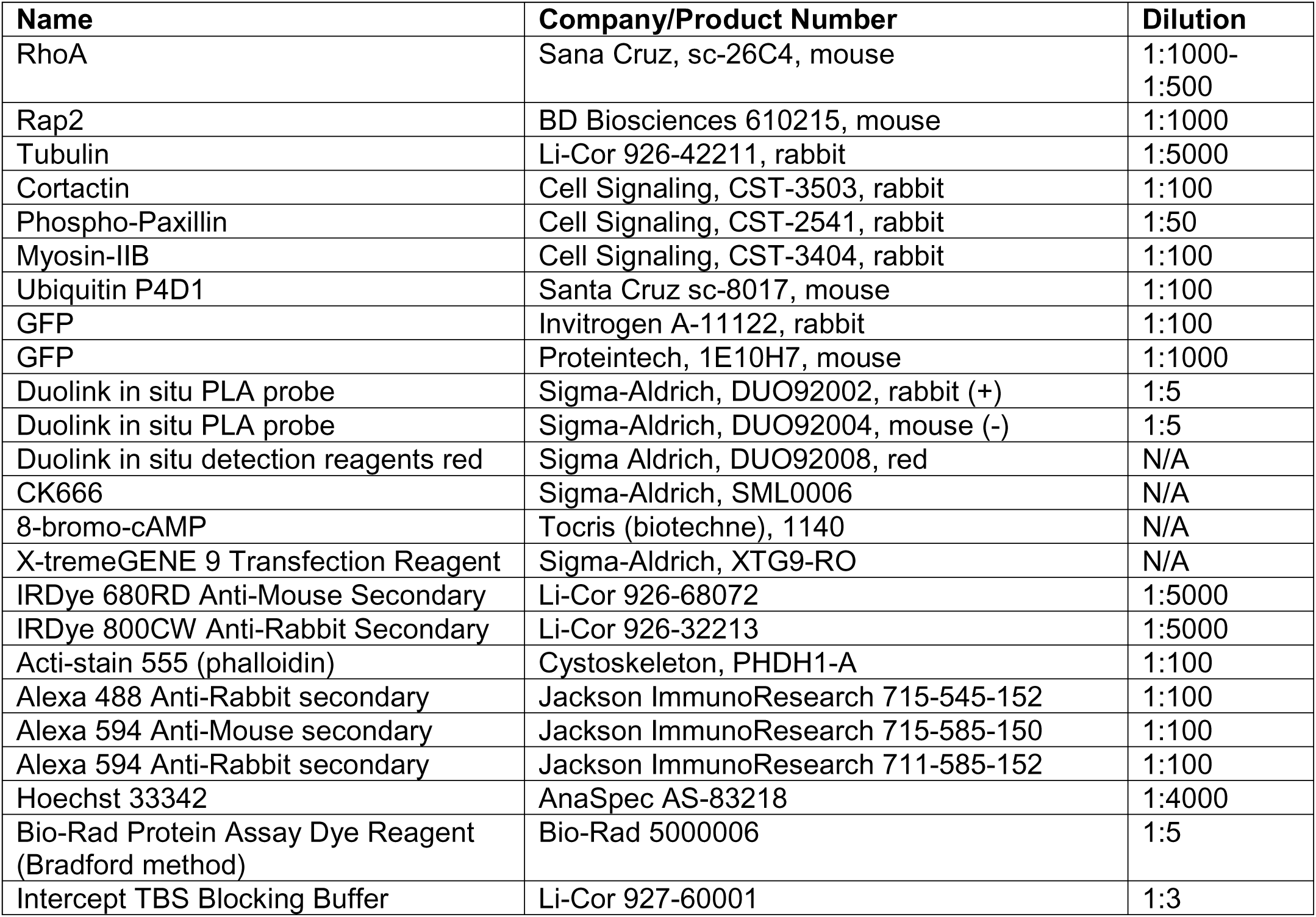
Antibodies and Reagents.

## SUPPLEMENTAL MATERIAL

Fig. S1 shows how data is calculated and further characterization of SACED as well as showing the effect of Rap2 KO on RhoA activity using whole cell lysate GST-RBD bead pulldown assays. Fig. S2 contains the controls used to validate PLA signal. Fig. S3 describes the failure of the Rap2-G12V mutant to localize abnormally or pulldown with GST-RalGDS beads. Fig. S4 shows how the Rap2-Q63E mutant effects actin ruffling. Fig. S5 contains the data of the CK666 washout experiment.

## ACKNOWLEDGEMENTS

This work was funded by the National Institutes of Health grant R01 GM122768 to R.P, grant S-MIP-22-60 from Research Council of Lithuania (to R.P.), and NIH T32 GM136444 to AN. The authors declare no competing financial interests. Thanks to Cecilia Caino and Raphael Garci-Matta for advice on RhoA activity pulldowns and to the Garcia-Matta lab for providing the dTom-2xrGBD RhoA biosensor. Thank you to Jeff Moore and Jeremy Brown for use of the spinning disk confocal microscope. Thank you to Paige Neumann for grammatical correction of this manuscript.

---

**Supplementary Figure 1:** Supplementary Figure 1 SACED analysis of the role of Rap2 on lamellipodia dynamics and Rap2 regulation of RhoA activity. **(A-D)** Kymographs taken from SACED time-lapse imaging of control and Rap2-KO cells (Videos 3 and 4). Annotated kymograph depicting the different measurements used for SACED calculations **(A).** Quantifications showing actin ruffle rate **(B)** distance of actin ruffle translocation **(C)** and membrane extension/retraction rates **(D). (D-E)** RhoA activity pulldown using GST-RBD beads which bind to active RhoA. Total RhoA pulled down was analyzed by western blot **(D)** and quantified **(E).**

**Supplementary Figure 2:** PLA Controls. **(A-C)** Control data from PLA experiments. Panel **(A)** shows PLA experiments performed with only an anti-GFP or an anti-ubiquitin antibody in cell with GFP-Rap2a-WT or GFP-Rap2a-K3R expression. Minimal PLA puncta are observed. Quantifications from PLA experiments including cell area **(B)** and the number of PLA puncta on the membrane across control and experimental conditions normalized per replicate to the median of the GFP-Rap2a-WT PLA data **(C).**

**Supplementary Figure 3:** The Rap2a-G12V mutation does not behave as a constitutively active mutation. **(A-B)** Phalloidin-596 (filamentous actin) staining of MDA-MB-231 cells expressing GFP-Rap2a-G12V or GFP-Rap2a-G12V-K3R. Quantification of GFP-Rap2a localization to the plasma membrane **(B). (C)** Rap2 activity assays performed using GST-RalGDS bead pulldown assay to examine the effect of the G12V mutation on Rap2a activation. Pulldown assay was performed using lysates from cells expressing GFP-Rap2a-G12V, GFP-Rap2a-G12V-K3R, or GFP-Rap2a-WT constructs and analyzed by western blot **(C)**.

**Supplementary Figure 4:** The Rap2a-Q63E mutation alters lamellipodia ruffling dynamics. **(A)** GFP-Rap2-Q63E expressing MDA-MB-231 cells were analyzed by time-lapse microscopy (see Video 9). Shown are selected GFP-Rap2a-Q63E and transmitted light images taken from time-lapse series. White arrow points to retracting ruffle with accumulating GFP-Rap2a-Q63E. **(B-D)** Kymographs generated from GFP-Rap2a-Q63E time-lapse series (Video 9). Quantification of GFP-Rap2a-Q63E ruffles dynamics as compared to GFP-Rap2a-WT ruffle dynamics (Fig. 1I, Video 5) showing changes in ruffle retraction rate **(C)** and translocation distance **(D)**.

**Supplementary Figure 5:** Activity is necessary for Rap2 retention at lamellipodia membrane. **(A-B)** MDA-MB-231 cells expressing either GFP-Rap2a-WT or GFP-Rap2a-K3R were treated with CK666 to inhibit macropinocytosis-dependent internalization of Rap2. CK666 was then washed out to resume the function of macropinocytic mechanisms. Quantification of Rap2 membrane localization is shown in **(B)**

## Notes

### Competing Interest Statement

The authors have declared no competing interest.

## REFERENCES

Bachmann, M., A. Skripka, K. Weißenbruch, B. Wehrle-Haller, and M. Bastmeyer. 2022. Phosphorylated paxillin and phosphorylated FAK constitute subregions within focal adhesions. J Cell Sci. 135.

Bisi, S., A. Disanza, C. Malinverno, E. Frittoli, A. Palamidessi, and G. Scita. 2013. Membrane and actin dynamics interplay at lamellipodia leading edge. Current Opinion in Cell Biology. 25:565–573.

Bos, J.L., H. Rehmann, and A. Wittinghofer. 2007. GEFs and GAPs: Critical Elements in the Control of Small G Proteins. Cell. 129:865–877.

Bravo-Cordero, J.J., L. Hodgson, and J. Condeelis. 2012. Directed cell invasion and migration during metastasis. Current Opinion in Cell Biology. 24:277–283.

Burridge, K., and C. Guilluy. 2016. Focal adhesions, stress fibers and mechanical tension. Exp Cell Res. 343:14–20.

Chen, W.T. 1979. Induction of spreading during fibroblast movement. Journal of Cell Biology. 81:684–691.

Choi, B.H., M.R. Philips, Y. Chen, L. Lu, and W. Dai. 2018. K-Ras Lys-42 is crucial for its signaling, cell migration, and invasion. J Biol Chem. 293:17574–17581.

Clague, M.J., and S. Urbé. 2017. Integration of cellular ubiquitin and membrane traffic systems: focus on deubiquitylases. Febs j. 284:1753–1766.

Collins, S.E., M.E. Wiegand, A.N. Werner, I.N. Brown, M.I. Mundo, D.J. Swango, G. Mouneimne, and P.G. Charest. 2023. Ras-mediated activation of mTORC2 promotes breast epithelial cell migration and invasion. Mol Biol Cell. 34:ar9.

Critchley, W.R., C. Pellet-Many, B. Ringham-Terry, M.A. Harrison, I.C. Zachary, and S. Ponnambalam. 2018. Receptor Tyrosine Kinase Ubiquitination and De-Ubiquitination in Signal Transduction and Receptor Trafficking. Cells. 7.

Dao, V.T., A.G. Dupuy, O. Gavet, E. Caron, and J. de Gunzburg. 2009. Dynamic changes in Rap1 activity are required for cell retraction and spreading during mitosis. J Cell Sci. 122:2996–3004.

Date, S.S., P. Xu, N.L. Hepowit, N.S. Diab, J. Best, B. Xie, J. Du, E.R. Strieter, L.P. Jackson, J.A. MacGurn, and T.R. Graham. 2022. Ubiquitination drives COPI priming and Golgi SNARE localization. Elife. 11.

Duncan, E.D., K.J. Han, M.A. Trout, and R. Prekeris. 2022. Ubiquitylation by Rab40b/Cul5 regulates Rap2 localization and activity during cell migration. J Cell Biol. 221.

Franz, C.M., G.E. Jones, and A.J. Ridley. 2002. Cell Migration in Development and Disease. Developmental Cell. 2:153–158.

Fregoso, F.E., T. van Eeuwen, G. Simanov, G. Rebowski, M. Boczkowska, A. Zimmet, A.M. Gautreau, and R. Dominguez. 2022. Molecular mechanism of Arp2/3 complex inhibition by Arpin. Nat Commun. 13:628.

Fuentes-Calvo, I., P. Crespo, E. Santos, J.M. López-Novoa, and C. Martínez-Salgado. 2013. The small GTPase N-Ras regulates extracellular matrix synthesis, proliferation and migration in fibroblasts. Biochim Biophys Acta. 1833:2734–2744.

Gopal Krishnan, P.D., E. Golden, E.A. Woodward, N.J. Pavlos, and P. Blancafort. 2020. Rab GTPases: Emerging Oncogenes and Tumor Suppressive Regulators for the Editing of Survival Pathways in Cancer. Cancers (Basel). 12.

Gorelik, R., and A. Gautreau. 2015. The Arp2/3 inhibitory protein arpin induces cell turning by pausing cell migration. Cytoskeleton. 72:362–371.

Guilluy, C., A.D. Dubash, and R. García-Mata. 2011. Analysis of RhoA and Rho GEF activity in whole cells and the cell nucleus. Nat Protoc. 6:2050–2060.

Hinz, B., W. Alt, C. Johnen, V. Herzog, and H.W. Kaiser. 1999. Quantifying lamella dynamics of cultured cells by SACED, a new computer-assisted motion analysis. Exp Cell Res. 251:234–243.

Hotulainen, P., and P. Lappalainen. 2006. Stress fibers are generated by two distinct actin assembly mechanisms in motile cells. Journal of Cell Biology. 173:383–394.

Janoueix-Lerosey, I., E. Pasheva, M.F. de Tand, A. Tavitian, and J. de Gunzburg. 1998. Identification of a specific effector of the small GTP-binding protein Rap2. Eur J Biochem. 252:290–298.

Julian, L., and M.F. Olson. 2014. Rho-associated coiled-coil containing kinases (ROCK). Small GTPases. 5:e29846.

Kaksonen, M., H.B. Peng, and H. Rauvala. 2000. Association of cortactin with dynamic actin in lamellipodia and on endosomal vesicles. J Cell Sci. 113 Pt 24:4421–4426.

Kolega, J. 2006. The Role of Myosin II Motor Activity in Distributing Myosin Asymmetrically and Coupling Protrusive Activity to Cell Translocation. Molecular Biology of the Cell. 17:4435–4445.

Kraynov, V.S., C. Chamberlain, G.M. Bokoch, M.A. Schwartz, S. Slabaugh, and K.M. Hahn. 2000. Localized Rac Activation Dynamics Visualized in Living Cells. Science. 290:333–337.

Kumar, N., P. Prasad, E. Jash, M. Saini, A. Husain, A. Goldman, and S. Sehrawat. 2018. Insights into exchange factor directly activated by cAMP (EPAC) as potential target for cancer treatment. Mol Cell Biochem. 447:77–92.

Kurokawa, K., R.E. Itoh, H. Yoshizaki, Y.O.T. Nakamura, and M. Matsuda. 2003. Coactivation of Rac1 and Cdc42 at Lamellipodia and Membrane Ruffles Induced by Epidermal Growth Factor. Molecular Biology of the Cell. 15:1003–1010.

Kurokawa, K., and M. Matsuda. 2005. Localized RhoA Activation as a Requirement for the Induction of Membrane Ruffling. Molecular Biology of the Cell. 16:4294–4303.

Langemeyer, L., R. Nunes Bastos, Y. Cai, A. Itzen, K.M. Reinisch, and F.A. Barr. 2014. Diversity and plasticity in Rab GTPase nucleotide release mechanism has consequences for Rab activation and inactivation. eLife. 3:e01623.

Lerosey, I., P. Chardin, J. de Gunzburg, and A. Tavitian. 1991. The product of the rap2 gene, member of the ras superfamily. Biochemical characterization and site-directed mutagenesis. J Biol Chem. 266:4315–4321.

Linklater, E.S., E.D. Duncan, K.J. Han, A. Kaupinis, M. Valius, T.R. Lyons, and R. Prekeris. 2021. Rab40-Cullin5 complex regulates EPLIN and actin cytoskeleton dynamics during cell migration. J Cell Biol. 220.

Mahlandt, E.K., J.J.G. Arts, W.J. van der Meer, F.H. van der Linden, S. Tol, J.D. van Buul, T.W.J. Gadella, and J. Goedhart. 2021. Visualizing endogenous Rho activity with an improved localization-based, genetically encoded biosensor. J Cell Sci. 134.

Mishra, Y.G., and B. Manavathi. 2021. Focal adhesion dynamics in cellular function and disease. Cellular Signalling. 85:110046.

Molinie, N., and A. Gautreau. 2017. The Arp2/3 Regulatory System and Its Deregulation in Cancer. Physiological Reviews. 98:215–238.

Mosaddeghzadeh, N., and M.R. Ahmadian. 2021. The RHO Family GTPases: Mechanisms of Regulation and Signaling. Cells. 10.

Mullins, R.D., J.A. Heuser, and T.D. Pollard. 1998. The interaction of Arp2/3 complex with actin: Nucleation, high affinity pointed end capping, and formation of branching networks of filaments. Proceedings of the National Academy of Sciences. 95:6181–6186.

Mylvaganam, S., S.A. Freeman, and S. Grinstein. 2021. The cytoskeleton in phagocytosis and macropinocytosis. Curr Biol. 31:R619–r632.

Naumanen, P., P. Lappalainen, and P. Hotulainen. 2008. Mechanisms of actin stress fibre assembly. Journal of Microscopy. 231:446–454.

Neumann, A.J., and R. Prekeris. 2023. A Rab-bit hole: Rab40 GTPases as new regulators of the actin cytoskeleton and cell migration. Frontiers in Cell and Developmental Biology. 11.

O’Connor, K., and M. Chen. 2013. Dynamic functions of RhoA in tumor cell migration and invasion. Small GTPases. 4:141–147.

O’Connor, K.L., B.-K. Nguyen, and A.M. Mercurio. 2000. Rhoa Function in Lamellae Formation and Migration Is Regulated by the α6β4 Integrin and Camp Metabolism. Journal of Cell Biology. 148:253–258.

Pannekoek, W.-J., J.R. Linnemann, P.M. Brouwer, J.L. Bos, and H. Rehmann. 2013. Rap1 and Rap2 Antagonistically Control Endothelial Barrier Resistance. PLOS ONE. 8:e57903.

Post, A., W.J. Pannekoek, B. Ponsioen, M.J. Vliem, and J.L. Bos. 2015. Rap1 Spatially Controls ArhGAP29 To Inhibit Rho Signaling during Endothelial Barrier Regulation. Mol Cell Biol. 35:2495–2502.

Prior, I.A., P.D. Lewis, and C. Mattos. 2012. A comprehensive survey of Ras mutations in cancer. Cancer Res. 72:2457–2467.

Reedquist, K.A., E. Ross, E.A. Koop, R.M. Wolthuis, F.J. Zwartkruis, Y. van Kooyk, M. Salmon, C.D. Buckley, and J.L. Bos. 2000. The small GTPase, Rap1, mediates CD31-induced integrin adhesion. J Cell Biol. 148:1151–1158.

Richter, S., R. Martin, H.O. Gutzeit, and H.-J. Knölker. 2021. In vitro and in vivo effects of inhibitors on actin and myosin. Bioorganic & Medicinal Chemistry. 30:115928.

Rothenberg, K.E., Y. Chen, J.A. McDonald, and R. Fernandez-Gonzalez. 2023. Rap1 coordinates cell-cell adhesion and cytoskeletal reorganization to drive collective cell migration in vivo. Curr Biol. 33:2587–2601.e2585.

Sartre, C., F. Peurois, M. Ley, M.H. Kryszke, W. Zhang, D. Courilleau, R. Fischmeister, Y. Ambroise, M. Zeghouf, S. Cianferani, Y. Ferrandez, and J. Cherfils. 2023. Membranes prime the RapGEF EPAC1 to transduce cAMP signaling. Nat Commun. 14:4157.

Sasaki, A.T., A. Carracedo, J.W. Locasale, D. Anastasiou, K. Takeuchi, E.R. Kahoud, S. Haviv, J.M. Asara, P.P. Pandolfi, and L.C. Cantley. 2011. Ubiquitination of K-Ras enhances activation and facilitates binding to select downstream effectors. Sci Signal. 4:ra13.

Schaks, M., G. Giannone, and K. Rottner. 2019. Actin dynamics in cell migration. Essays in Biochemistry. 63:483–495.

Segev, N. 2011. GTPases in intracellular trafficking: an overview. Semin Cell Dev Biol. 22:1–2.

Simanov, G., I. Dang, A.I. Fokin, K. Oguievetskaia, V. Campanacci, J. Cherfils, and A.M. Gautreau. 2021. Arpin Regulates Migration Persistence by Interacting with Both Tankyrases and the Arp2/3 Complex. Int J Mol Sci. 22.

Suraneni, P., B. Rubinstein, J.R. Unruh, M. Durnin, D. Hanein, and R. Li. 2012. The Arp2/3 complex is required for lamellipodia extension and directional fibroblast cell migration. Journal of Cell Biology. 197:239–251.

Tanner, N., O. Kleifeld, I. Nachman, and G. Prag. 2019. Remodeling Membrane Binding by Mono-Ubiquitylation. Biomolecules. 9.

Trepat, X., Z. Chen, and K. Jacobson. 2012. Cell migration. Compr Physiol. 2:2369–2392.

Vetter, I.R., and A. Wittinghofer. 2001. The guanine nucleotide-binding switch in three dimensions. Science. 294:1299–1304.

Wang, Y., and J. Adjaye. 2011. A cyclic AMP analog, 8-Br-cAMP, enhances the induction of pluripotency in human fibroblast cells. Stem Cell Rev Rep. 7:331–341.

Watanabe, N., T. Kato, A. Fujita, T. Ishizaki, and S. Narumiya. 1999. Cooperation between mDia1 and ROCK in Rho-induced actin reorganization. Nature Cell Biology. 1:136–143.

Weed, S.A., A.V. Karginov, D.A. Schafer, A.M. Weaver, A.W. Kinley, J.A. Cooper, and J.T. Parsons. 2000. Cortactin localization to sites of actin assembly in lamellipodia requires interactions with F-actin and the Arp2/3 complex. J Cell Biol. 151:29–40.

Wehrle-Haller, B. 2012. Structure and function of focal adhesions. Current Opinion in Cell Biology. 24:116–124.

Weissman, A.M., N. Shabek, and A. Ciechanover. 2011. The predator becomes the prey: regulating the ubiquitin system by ubiquitylation and degradation. Nat Rev Mol Cell Biol. 12:605–620.

Wolfenson, H., A. Bershadsky, Y.I. Henis, and B. Geiger. 2011. Actomyosin-generated tension controls the molecular kinetics of focal adhesions. Journal of Cell Science. 124:1425–1432.

Wong, D.C.P., C.Q. Pan, S.Y. Er, T. Thivakar, T.Z.Y. Rachel, S.H. Seah, P.J. Chua, T. Jiang, T.W. Chew, P.K. Chaudhuri, S. Mukherjee, A. Salim, T.A. Aye, C.G. Koh, C.T. Lim, P.H. Tan, B.H. Bay, A.J. Ridley, and B.C. Low. 2023. The scaffold RhoGAP protein ARHGAP8/BPGAP1 synchronizes Rac and Rho signaling to facilitate cell migration. Molecular Biology of the Cell. 34:ar13.

Xu, P., H.M. Hankins, C. MacDonald, S.J. Erlinger, M.N. Frazier, N.S. Diab, R.C. Piper, L.P. Jackson, J.A. MacGurn, and T.R. Graham. 2017. COPI mediates recycling of an exocytic SNARE by recognition of a ubiquitin sorting signal. Elife. 6.

Zhang, H., Y.C. Chang, M.L. Brennan, and J. Wu. 2014. The structure of Rap1 in complex with RIAM reveals specificity determinants and recruitment mechanism. J Mol Cell Biol. 6:128–139.

