## Supplementary figures and images for "Mono-Ubiquitylation-Dependent Rap2 Activation Regulates Lamellipodia Dynamics During Cell Migration"

### Supplemental Figure 1

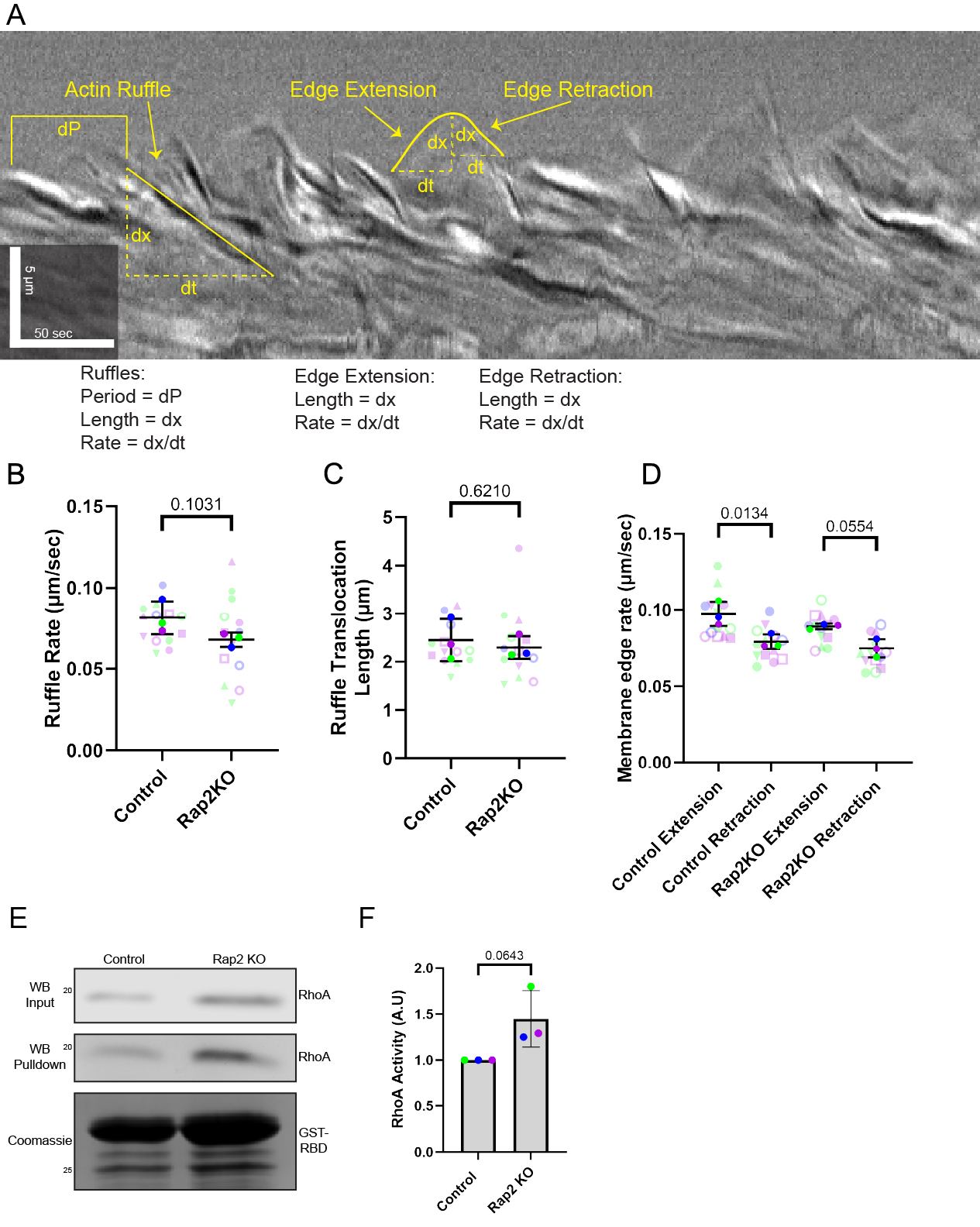

### Supplemental Figure 2

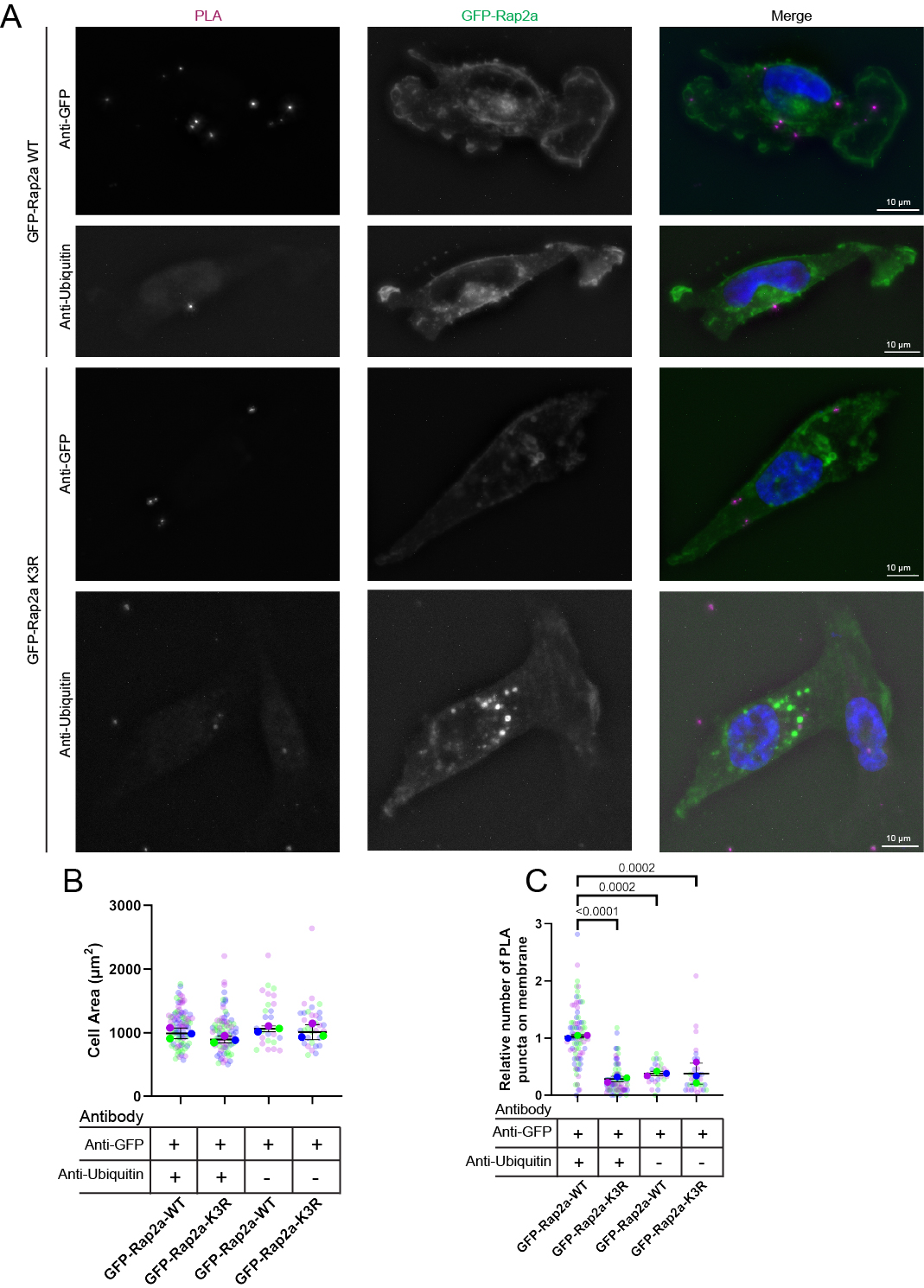

### Supplemental Figure 3

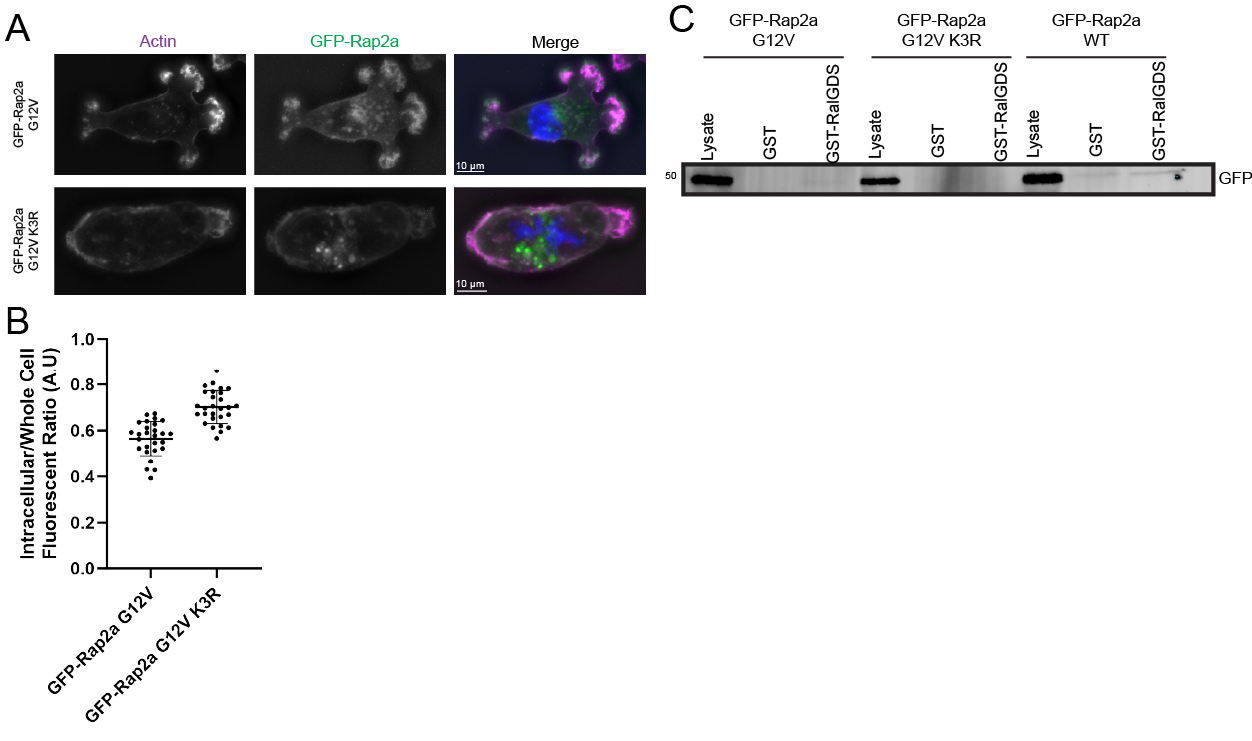

### Supplemental Figure 4

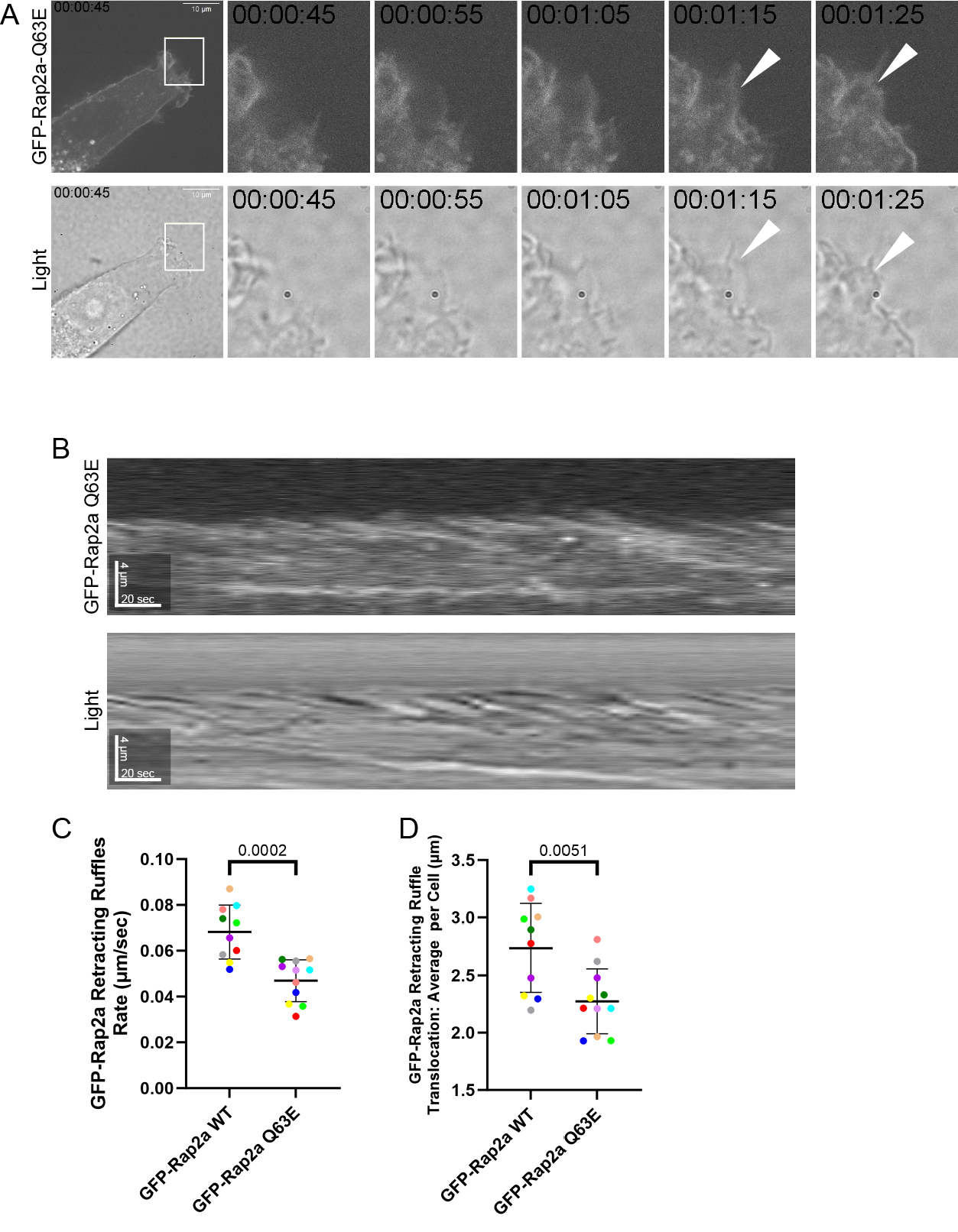

### Supplemental Figure 5

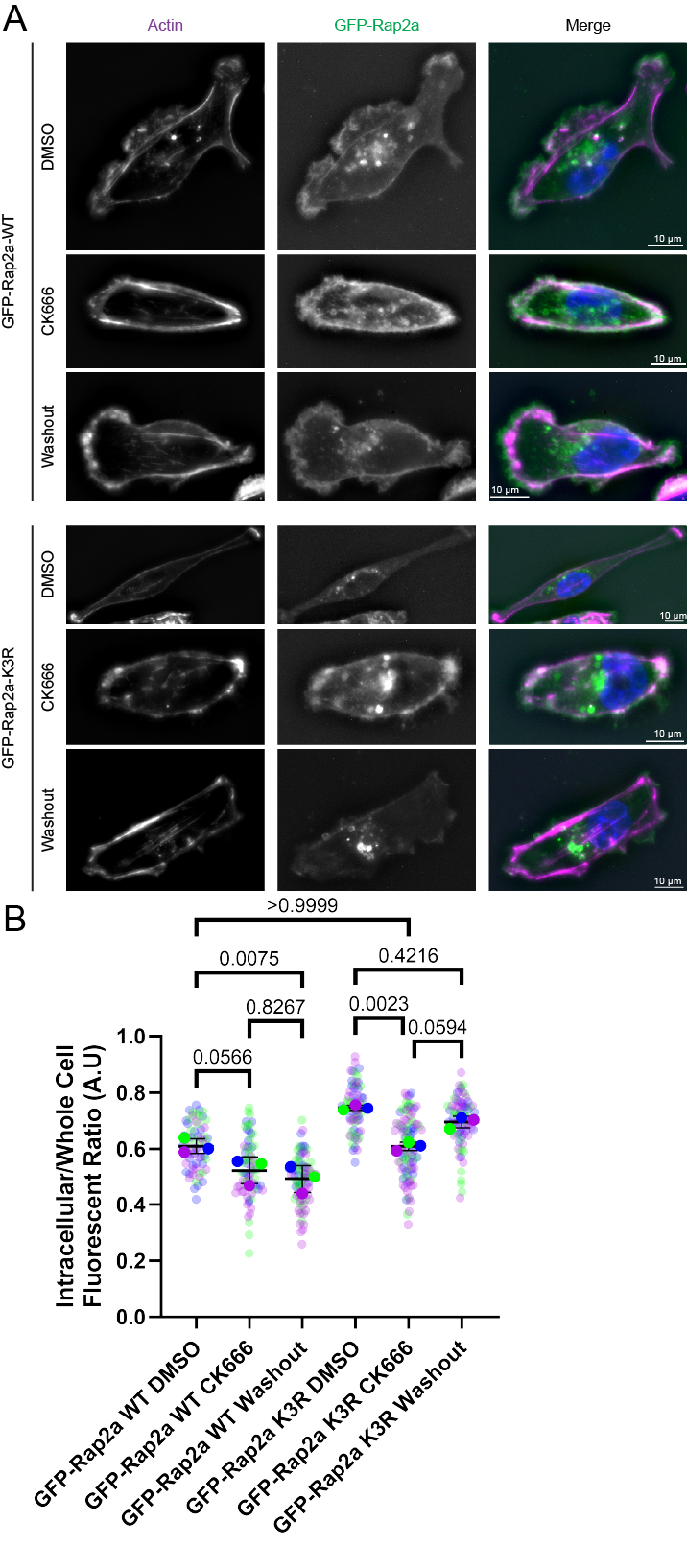
